# Probabilistic machine learning ensures accurate ambient denoising in droplet-based single-cell omics

**DOI:** 10.1101/2022.01.14.476312

**Authors:** Caibin Sheng, Rui Lopes, Gang Li, Sven Schuierer, Annick Waldt, Rachel Cuttat, Slavica Dimitrieva, Audrey Kauffmann, Eric Durand, Giorgio G. Galli, Guglielmo Roma, Antoine de Weck

**Affiliations:** Disease area Oncology, Novartis Institute for Biomedical Research, CH-4002 Basel, Switzerland; Chemical Biology and Therapeutics, Novartis Institute for Biomedical Research, CH-4002 Basel, Switzerland

## Abstract

Droplet-based single-cell omics, including single-cell RNA sequencing (scRNAseq), single-cell CRISPR perturbations (e.g., CROP-seq), and single-cell protein and transcriptomic profiling (CITE-seq) hold great promise for comprehensive cell profiling and genetic screening at the single-cell resolution. However, these technologies suffer from substantial noise, among which ambient signals present in the cell suspension may be the predominant source. Current models to address this issue are highly technology-specific and relatively scRNAseq-centric. while a universal model to describe the noise across these technologies may reveal this common source, improving the denoising accuracy. To this end, we explicitly examined these unexpected signals in multiple datasets across droplet-based technologies, summarised a predictable pattern, and developed single-cell Ambient Remover (scAR) – a hypothesis-driven machine learning model to predict and remove ambient signals (including mRNA counts, protein counts, and sgRNA counts) at the molecular level. We benchmarked scAR on three technologies – single-cell CRISPR screens, CITE-seq, and scRNAseq along with the state-of-the-art single-technology-specific approaches. scAR showed high denoising accuracy for each type of dataset.

## Introduction

Single-cell RNA sequencing (scRNAseq) enables researchers to investigate transcriptomes at single-cell resolution, improving our understanding of cellular heterogeneity and interactions between single cells and the microenvironment. Recent efforts have extended scRNAseq beyond transcriptomes by encoding additional layers of information, resulting in versatile tools for single-cell omics. For instance, by combining functional screens with scRNAseq, single-cell CRISPR screens have enabled the interrogation of multiple biological nodes in a single experiment^1–3^. By combining ssDNA-barcoded-antibodies with scRNAseq, CITE-seq has simultaneously quantified mRNA and surface proteins for immunophenotyping^4^. The most recent effort has combined both technologies to enable multimodal profiling of transcriptome and surface proteins in response to gene perturbations in cancer cells^5^.

Applying these pioneering technologies is challenging despite the exciting concepts and anticipated potential. One outstanding reason is the vast presence of measurement noise. Various technical factors, such as ambient contamination^6,7^, amplification bias^8^, and index swapping^9^ generate noise in single-cell omics experiments. Many methods have been proposed to deal with the background signals and have facilitated the application of single-cell technologies^6,7,10–12^. Most of them are specific to transcriptome data^6,7,10,11^, and several specialise in cleaning protein data in CITE-seq^12,13^, while few focus on recovering identity barcodes from noisy observations in single-cell feature-barcoded technologies (such as single-cell CRISPR screens and cell indexing^14–16^). These approaches are generally not transferrable between technologies. However, conceptually, the mentioned single-cell technologies have little technical difference in capturing the respective molecules (i.e., mRNA, sgRNA, expressed barcodes, and antibodies). All relevant experimental objects (cells and molecules) undergo similar processes, such as droplet formation, cell lysis, library construction, and sequencing. Background noise likely originates in a similar (if not identical) way in each of these technologies, meaning an ideal model can, in principle, describe the common sources of the artefacts in a non-technology-specific manner. Unfortunately, no such algorithm has been proposed so far.

Here, we took a bottom-up strategy – investigating ambient signals across technologies, summarising a predictable pattern, and building a statistical deep learning model, termed single cell ambient remover (scAR) to model the common noise sources. We first evaluated the potential noise source across technologies. Cell-free transcripts have been observed in empty droplets^17^. They may arise from ambient mRNAs in single-cell suspension, which likely originates from damaged cells (e.g., caused by cell lysis)^6,18,19^. This hypothesis suggests that ambient mRNAs may not be completely random but deterministic signals to a certain extent. Indeed, the frequencies of cell-free transcripts correlate well with their cellular counterparts^6^. These motivated us to systematically evaluate the ambient signal hypothesis in multiple single-cell technologies, which revealed a predictable pattern of ambient noise. Exiting single-cell analysis methods which implement probabilistic machine learning technologies, including totalVI^12^, DCA^10^, scVI^11^, DecontX^7^, and scVAE^20^, have shown significant advantages in modelling single-cell omics data. Together with the ambient signal hypothesis, these inspired us to build a universal probabilistic model to describe this type of noise across technologies. To highlight the generality of our approach, we applied scAR to multiple kinds of single-cell datasets – including an internal single-cell CRISPR screens dataset and several public CITE-seq datasets and scRNAseq datasets – and demonstrated scAR’s accuracy in comparison with competing technology-specific methods where available.

## Results

### The scAR model

scAR uses a latent variable model to represent the biological and technical components in the observed count data (Fig. 1). It is designed under the ambient signal hypothesis, which assumes that ambient signals originate from broken cells during sample preparation, are homogenised in the single-cell solution (ambient signal pool) and contaminate cell-containing and cell-free droplets (Fig. 1a). Mathematically speaking, we assume that ambient signals in cell-containing and cell-free droplets are drawn from a distribution (Binomial, or Poisson, or zero-inflated Poisson) with fixed probabilities per feature (denoted as ambient frequencies, α). These probabilities can be estimated by averaging cell-free droplets as their counts all are ambient signals. For cell-containing droplets, we introduce two hidden variables, noise ratios (ε) and native expression frequencies (β), to represent the total contamination level per cell and normalised ‘true’ expression per cell, respectively (Fig. 1b). scAR simultaneously infers ε and β using the variational autoencoder (VAE) framework^21–23^ (Fig. 1b, Methods).

**Fig. 1.**
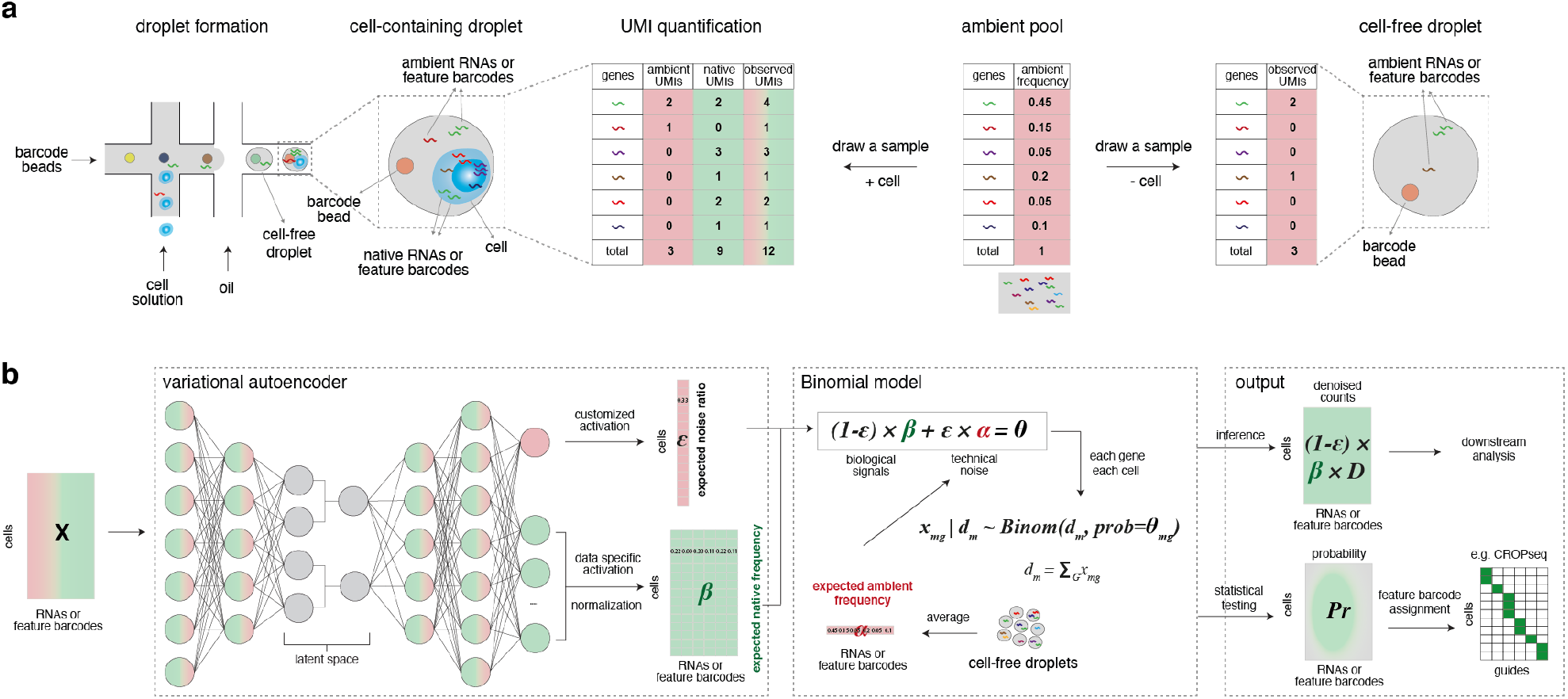
The overview of scAR. scAR is a hypothesis-driven denoising model for droplet-based single-cell omics technologies. **a**, Demonstration of the ambient signal hypothesis. During the preparation of the single-cell solution, RNA or protein counts are released upon cell lysis and consequently encapsulated by droplets with or without a cell. In the case of a cell, these exogenous molecules are mixed with native ones and barcoded by the same 10x bead, resulting in ravelling count data. Under this assumption, the ambient signals in both cell-containing and cell-free droplets are drawn from the same pool. UMI stands for the unique molecular identifier. For illustration, the reddish-purple, the light green, and the mixture indicate ambient signals, native signals, and observed counts, respectively. **b**, scAR takes raw count matrices of RNA or protein as input and learns two sets of parameters (ε and β) through the variational autoencoder. ε, a column vector represents noise ratios per cell and β, a matrix represents cell-wise native frequencies of RNAs or proteins. α, a row vector represents the ambient frequencies of RNAs or proteins, which is empirically estimated by averaging cell-free droplets. scAR uses a unique α for all cells in a single experiment. The observed raw counts are modelled using a Binomial model containing known parameters α and sequencing depth D and two hidden variables ε and β. We optimised ε and β by minimising the reconstruction errors and K-L divergence (Methods). scAR outputs two matrices, a denoised count matrix and a matrix of Bayes factors. The latter represents the likelihood that native signals are present in given observed counts. The meaning of colour codes is the same as **a**.

We use the optimised variables ε and β as well as the sequencing depth to estimate the ‘theoretical’ gene expression, which is denoted as denoised counts and can be directly used by other well-established tools for downstream analysis. In addition, in several feature barcode technologies, such as CROP-seq and CellTagging^14^, the presence/absence of native signals is more critical information than the actual level. To reflect this, scAR also outputs a matrix of Bayes factors representing the likelihood of whether ‘true’ native signals are present in raw counts.

### Evaluation of ambient signal hypothesis

We conducted a case study that combined CROP-seq and bulk sequencing to evaluate the ambient signal hypothesis (Fig. 2a and Methods). We designed a viral pool of 99 sgRNAs targeting 13 different genes (supplementary table I), most of them being essential in MCF7 cells^24,25^ (supplementary Fig. 1a). We infected MCF7 cells expressing dCas9-KRAB with the lentiviral libraries at 0.3 multiplicity of infection to ensure a majority (~84% after selection) of single infections. Excessive sgRNAs in a cell are supposed to be ambient signals. Cells were harvested at various time points post-transduction and split into two portions, with one portion taken for 10x scRNAseq and the other for bulk sequencing of sgRNAs, which revealed the frequencies of sgRNA libraries in the samples. In addition, the downregulation of target genes by CRISPRi provided an additional way to assess the identification of truly integrated sgRNAs.

**Fig. 2.**
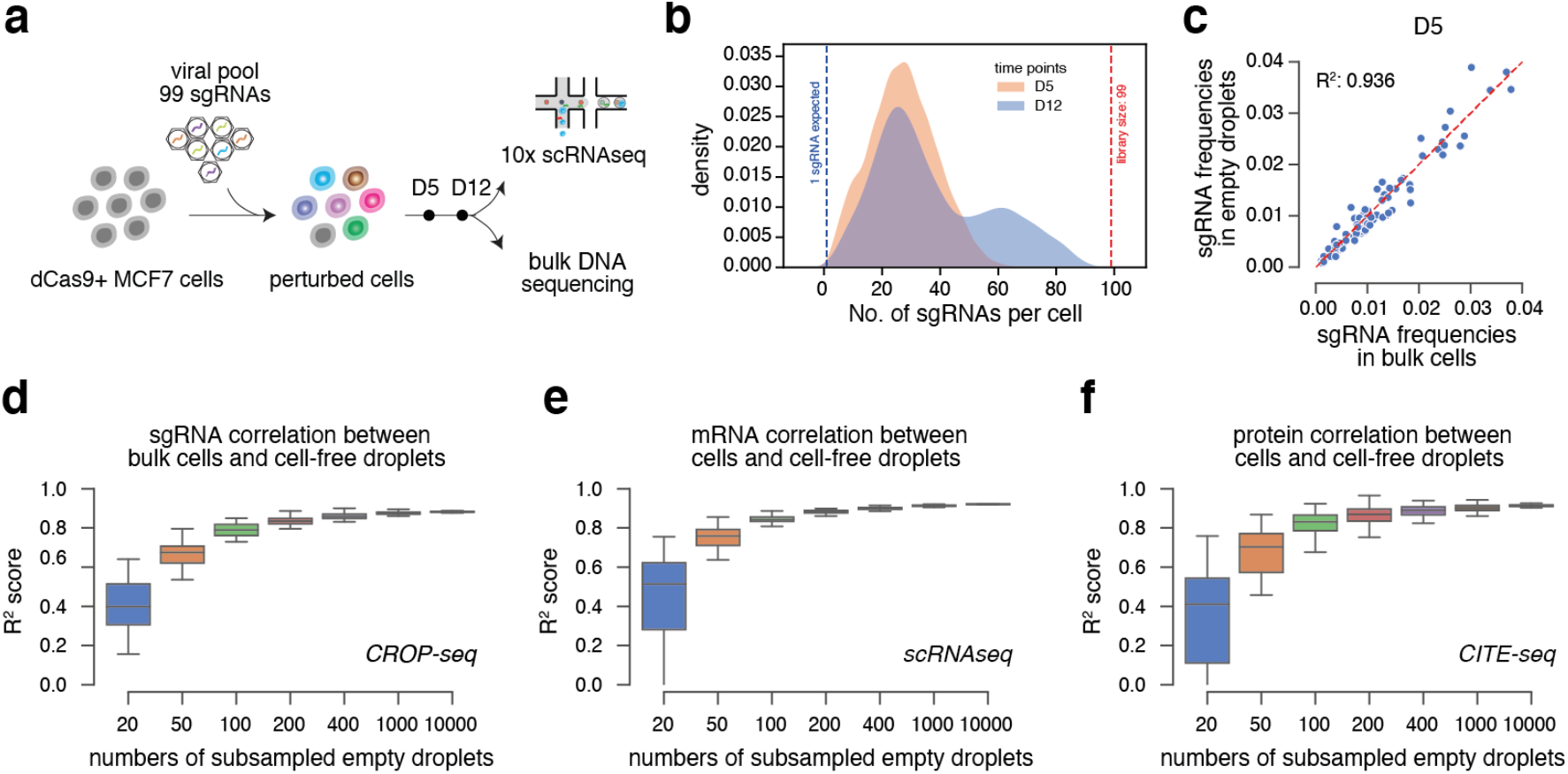
Evaluation of the ambient signal hypothesis. We systematically evaluated the ambient signal hypothesis in various types of single-cell omics data. **a**, Demonstration of the experimental design (Methods). **b**, Distribution of distinct sgRNAs per cell in the raw counts. The X-axis represents the number (not UMI count) of distinct sgRNAs per cell; the y-axis shows the density. The size of lentivirus libraries and the expected number of sgRNA are highlighted by dashed lines. Time points were indicated. **c**, Scatterplot of sgRNA frequencies between two independent experiments. The X-axis represents sgRNA frequencies in bulk DNA sequencing; Y-axis represents sgRNA frequencies obtained by averaging RNA counts in cell-free droplets from 10x scRNAseq. Each dot represents an sgRNA, the red dashed lines represent y=x, and coefficients of determination (*R*^2^ scores) is shown. **d**, Coefficients of determination (*R*^2^ scores) show the correlation of sgRNA frequencies between bulk cells and randomly sampled cell-free droplets. The X-axis shows the numbers of randomly sampled cell-free droplets. Forty samplings were performed in each group. **e-f**, Similar to **Fig. 2d**, correlation analysis of mRNA counts and protein counts (**f**) between cell-containing and -free droplets.

We first examined the raw sgRNA counts in cells and observed the strong presence of ambient counts (Fig. 2b). ~25 distinct sgRNAs were detected per cell on average, while <=1 sgRNA was expected because of the low multiplicity of infection. To validate whether ambient sgRNAs are correlated with their native counterparts, we compared sgRNA frequencies in bulk sequencing and cell-free droplets from CROP-seq. Results showed a high correlation of sgRNA frequencies at both time points (Fig. 2c and supplementary Fig. 1b). Randomly sampled subsets of cell-free droplets (from 20 to 10000) show a high correlation (Fig. 2d). Together, these observations confirm that ambient signals are not random noise but endogenous expression-correlated artefacts. We next asked whether this correlation persists in other droplet-based technologies. To answer it, we checked the mRNA counts of the in-house CROP-seq experiment, protein counts of a public CITE-seq^26^, and sgRNA counts of another three single-cell CRISPR screens^3,26,27^. Surprisingly, feature frequencies (i.e., mRNA, sgRNA, or protein) in as few as ~50 cell-free droplets consistently show a high correlation to that of the cell-containing droplets despite the varying numbers (from 32~20,000) of features (Fig. 2e-f and supplementary Fig. 2). Altogether, these indicate that the ambient signals are systematic noise across single-cell omics technologies, and they are strongly correlated with native signals in the pooled cells. Thus, building scAR on the ambient hypothesis is rational.

### scAR enables accurate guide calling in single-cell CRISPR screens

Single-cell CRISPR screens (including Perturb-seq^28^, CRISP-seq^29^, and CROP-seq^1,2^) are powerful tools for functional screening with single-cell transcriptome readout, yet, few methods focused on dealing with the noise, although accurate guide calling is a key to data quality and result interpretation. Most of the current studies considered the most highly expressed guides as the true signal and relatively low counts as background noise^1,2,28–30^, which we found inappropriate, especially when a few guides were predominantly enriched in the libraries (data not shown), as these guides could cause a significant amount of ambient signal and mask the true native signals.

We, therefore, asked whether scAR could distinguish the ambient signals and improve the guide calling. To test it, we applied scAR to the CROP-seq dataset and benchmarked it against the baseline approach, termed as naïve assignment. Here, we did not involve any subjective filters to either naïve assignment or scAR-based assignment for the benchmarking purpose. All cells that passed the default gene and cell filtering in Cellranger were included for downstream analysis (Methods). By naïve assignment, ~80% cells (20076 out of 25248, D5 and D12 combined) were assigned to unique guides, and ~20% (5170 out of 25248, D5 and D12 combined) cells were assigned to multiple (>=2) guides due to equal expression (Fig. 3a). ‘Multiple-infected’ cells were generally filtered out before downstream analysis in CROP-seq experiments; in other words, naïve assignment caused loss of ~20% cells. scAR estimated the expected ambient counts then compared them to the observed counts via hypothesis testing to evaluate the probability of the presence of native signals. It nearly assigned all cells to a single guide (Fig. 3a) despite ~20% cells with equally expressed guides. We next examined the cell fraction partitioned by distinct sgRNAs after guide assignment. This fraction was expected to be identical to sgRNA frequencies in bulk sequencing since both reflected sgRNA libraries in the cell pool. scAR-resulting cell fractions were highly correlated with sgRNA frequencies in bulk sequencing (*R*^2^=0.879 at D5 and *R*^2^=0.910 at D12), while naïve assignment preferably over-assigned a few sgRNAs of highest expression (*R*^2^=0.369 at D5 and *R*^2^=0.861 at D12, Fig. 3b-c).

**Fig. 3.**
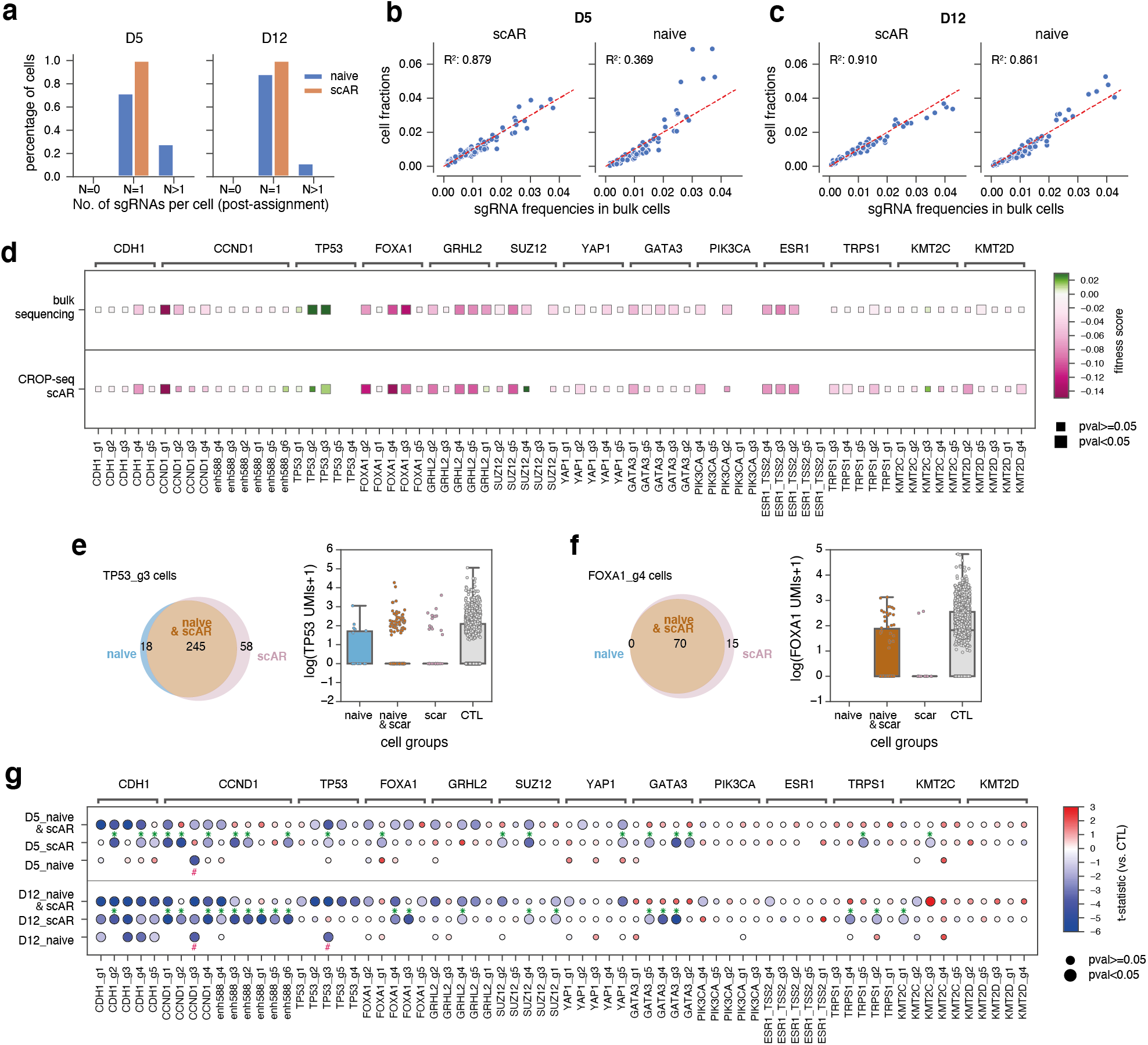
scAR enables accurate guide calling in single-cell CRISPR screens. We demonstrate that scAR-guided assignment shows superior performance compared to the widely used naïve assignment. **a**, Distribution of null, single and multiple assignments. Cells were grouped by the number of assigned sgRNA(s). **b-c**, Scatterplots show the correlation of sgRNA frequencies between cells from 10x scRNAseq (post-assignment) and bulk cells at D5 (**b**) and D12 (**c**). Y-axis represents cell fractions grouped by distinct sgRNAs. The red dashed lines represent y=x. **d**, CROP-seq with scAR-guided assignment shows comparable results of gene dependency analysis to bulk screening. Green and maroon indicate cancer cell promotion and cancer growth inhibition, respectively. Sizes of squares indicate p values. **e-f**, Two selected guides demonstrate scAR’s sensitivity and specificity. The Venn diagrams show the number of cells assigned with TP53_g3 (**e**) at D5 and FOXA1_g4 (**f**) at D12. Exclusive naïve assignment, exclusive scAR assignment, and accordant assignment are marked with blue, purple, and brown, respectively. The boxplots show the expression of TP53 or FOXA1 in these subgroups. Y-axis represents the log-transformed UMI counts after library size normalisation (Methods). Dots represent cells. Co-assigned CTL cells are used as the negative control. **g**, The dotplot shows the on-target downregulation. The X-axis represents guide groups. Y-axis represents subgroups of cells, separated by two time points and assignment approaches. Target genes are shown on the top. Their expression in each group is compared with that in the CTL group (centred at zero), and resulting t-statistics are guided by the dot colours (Methods, see t-statics of z-normalized expression in supplementary Fig. 4a). Blue and red indicate down- and up-regulation, respectively. The bimodal sizes of circles represent the p-values from the t-test. The asterisks (*) and hashes (#) highlight the guide groups where scAR outperforms and underperforms naïve assignment, respectively.

Pooled screens with proliferation readouts led to the discovery of many potential drug targets in Oncology^24,25,31^, where the gene dependencies were generally calculated on millions of reads per target^32^. We wondered whether a few hundred cells per guide in the CROP-seq experiment could capture the gene dependencies given the high accuracy of guide assignment by scAR. To check it, we first used bulk sequencing data of multiple time points (D0, D5, D7, D9, and D12) to fit an exponential model for each guide (see our previous report^3^). The growth rate in the model reflected the fitness score of each guide, which were consistent with well-established datasets (supplementary Fig. 1a). We then took the same procedure but with the cell fraction data (CROP-seq experiment) of three time points (D0, D5, and D12) and compared the resulting dependency scores (Fig. 3d). Excitingly, most of the guides show comparable fitness scores between these two datasets, despite the small sample sizes (~92 cells per guide) and limited time points. This confirms the high accuracy of scAR-based assignments.

Next, we checked the expression levels of targeted genes in cells with certain guides assigned exclusively by either naïve approach or scAR (Fig. 3e-g). We considered the cells assigned by both naïve and scAR as the positive control and cells assigned with CTL sgRNAs by both naïve and scAR as the negative control. In an example shown in Fig. 3e, naïve assignment assigned 263 cells to TP53_g3, 245 cells among which were mutually assigned to the same guide by scAR. These 245 cells show downregulation of TP53, suggesting the effectiveness of this guide. However, the remaining naïve assigned 18 cells show similar expression as in CTL cells, suggesting that these cells may not integrate TP53_g3. More importantly, another 58 TP53_g3 cells, exclusively identified by scAR, show low expression as in the mutually assigned cells. Similarly, for the other example, FOXA1_g4 (Fig. 3f), 15 cells identified by scAR but missed by naïve assignment show a similar expression pattern as the mutually assigned cells. To systematically assess and visualise the difference, we performed a t-test on the expression of targets among these subgroups for each guide. We visualized both t-statics and p-values using dotplots (Fig. 3g and supplementary Fig. 3a). In total, scAR rescued ~20 sgRNA groups at each time point which were missed by naïve assignment as confirmed by statical confidences. In comparison, fewer than two sgRNA groups were missed by scAR at each time point, compared to naïve assignment. In addition, we counted the cell numbers of each subgroup and visualised the difference (supplementary Fig. 3b). Naïve assignment over-assigned cells to a few guides, such as CCND1_g3, CTL_g5, and YAP1_g1, likely due to their more substantial ambient presence than other guides (supplementary Fig. 3c), whereas the power of scAR to identify ambient sgRNAs by their distribution ensured unbiased assignment. It recovered more than ten cells per group for most of the target groups (supplementary Fig. 3b). Together, by inspecting the guide assignment in the CROP-seq dataset, we showed that scAR significantly improved the assignment accuracy in feature barcode technologies, where the presence rather than the quantity of native signals was the key information.

### scAR identifies a resistant marker

Next, we evaluated scAR on CITE-seq along with current protein denoising methods,

### scAR removes the ambient protein counts in CITE-seq

Next, we evaluated scAR on CITE-seq along with current protein denoising methods, such as totalVI^12^ and DSB^13^. We used a public CITE-seq dataset^26^ of peripheral blood mononuclear cells (PBMC5k), which were profiled with a panel of 32 antibody-conjugated oligos consisting of surface markers of B cells, T cells, Natural killer cells (NK), monocytes, and Dendritic cells (DC) (supplementary Fig. 4a). We first annotated cell types by clustering cells with transcriptome signatures (Fig. 4a and Methods) and compared protein counts per cell type before and after denoising.

**Fig. 4.**
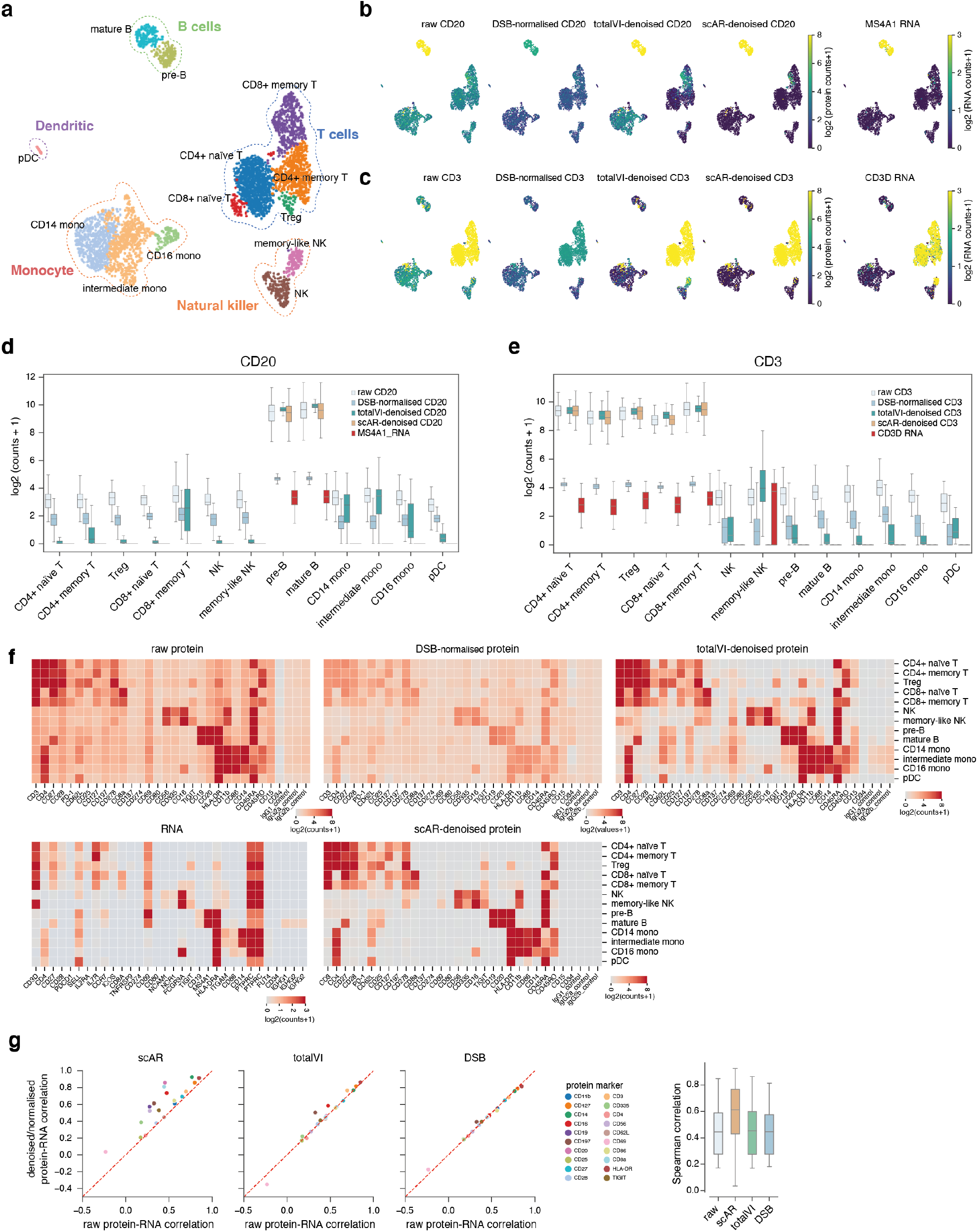
scAR removes the ambient protein counts in CITE-seq data. scAR outperforms competing protein-denoising approaches. **a**, UMAP of the PBMC5k dataset. Cell types are annotated using transcriptome data (Methods). **b-e**, Two selected examples, CD20 antibody (**b, d**) and PD-1 antibody (**c, e**), demonstrate scAR’s performance. UMAPs visualise raw protein counts, DSB-normalised, totalVI-denoised, and scAR-denoised protein counts, and the corresponding RNA counts, respectively. Colour bars represent log2 (counts +1) (log2(normalised values + 1) in the case of DSB), and UMAPs of protein counts use the same scaling. Boxplots show the single-cell counts grouped by cell types. **f**, Heatmaps show the cell types’ average protein (raw, DSB-normalised, totalVI-denoised, and scAR-denoised) and RNA counts. Columns and rows represent the antibodies (or corresponding RNAs) and cell types. **g**, Scatterplots show the Spearman correlation coefficients between RNA-protein pairs before (x-axis) and after (y-axis) protein denoising. The red dashed line represents y=x. Dots represent antibodies. The rightest boxplot summarises the coefficients.

As expected, ambient protein signals were observed in raw counts in every cell. Both scAR and totalVI could preserve cell-specific true signals, and scAR nearly eliminated all ambient counts, while totalVI generally missed some ambient counts in several cell types (especially monocytes and memory-like NK cells, Fig. 4b-e and supplementary Fig. 4). DSB-denoised values reflected the number of standard deviations above background noise rather than the molecular-level counts; it rescaled the true signals and hardly eliminated all ambient signals in cells given the high cell-to-cell variations (even for control antibodies, a half population was supposed to have positive values with DSB normalisation). For example, scAR cleaned most of the non-specific CD20 and CD19 in non-B cells, and totalVI also removed most of non-specific CD20 and CD19 but missed some in monocytes and memory-like NK cells. DSB rescaled both the native and ambient signals and could not clean all ambient signals in other cells (Fig. 4b, 4d, and supplementary Fig. 4b). For simplicity and fairness, we benchmark scAR with only totalVI when comparing the counts of proteins in the following text as both are molecular level denoising approaches. Similarly, we found that scAR nearly recognised all non-specific CD3 and PD-1 in non-T cells (Fig. 4c, 4e, and supplementary Fig. 4c), non-specific CD56 and CD335 in non-NK cells (supplementary Fig. 4d-e) as well as non-specific CD14 in non-monocytes (supplementary Fig. 4f). totalVI also showed promising performance, identifying all non-specific CD14 in non-monocytes (supplementary Fig. 4f), almost all non-specific CD56 and CD335 in non-NK cells (supplementary Fig. 4d-e) and most non-specific CD3 and PD-1 in non-T cells (Fig. 4c, 4e, and supplementary Fig. 4c). We also found that scAR successfully identified the IgG1 signal as noise in all cells while totalVI had a visible level of negative-positive rate (supplementary Fig. 4g). It is worth underlining that the expressions of ambient proteins were highly diversified (mean 9.6 ± STD 43.4 per marker per cell). In some cases, the true signals were as low as background noise. For example, the naïve T cells showed a similar PD-1 level as B cells, NK cells, and monocytes; while scAR retained the PD-1 signal in naïve T cells but eliminated all signals in B cells, NK cells, monocytes, and DC cells (supplementary Fig. 5a-b). To assess how much scAR corrected the ambient noise, we calculated the per cell fractions of estimated ambient counts and compared it with totalVI (supplementary Fig. 6). In addition to removing ambient counts, totalVI added extra counts, particularly to the highly expressed proteins due to its imputation functionality^12^, resulting in unchanged total counts and high denoising variance (noise ratio at −0.2% ± STD 29%). In contrast, scAR removed more raw protein counts on average but was highly consistent in each cell type (mean 7.5% ± STD 2.4%).

We next compared the averaged protein levels by cell type before and after denoising and aligned them with RNA expressions (Fig. 4f). Almost all antibodies showed intense background noise at the populational level. DSB-resulted normalisation down-scaled but did not eliminate ambient signals, and totalVI preserved true signals and reduced most ambient contamination. scAR also held true signals and nearly removed all the background noise, resulting in highly specific expressions of markers. For example, scAR-denoised CD8 were exclusively present in CD8+ T cells, while after DSB- or totalVI-denoising, there was still residual CD8 in other cell types (Fig. 4f and supplementary Fig. 7a). Compared to DSB and totalVI, scAR-denoised CD197 and CD45RA could separate Naïve and memory T cells well (Fig. 4f and supplementary Fig. 7b). Overall, after scAR-denoising, protein expressions correlated stronger with corresponding RNAs, and the Spearman correlation coefficient between them was increased to ~0.6 (Fig. 4g). Together, these results demonstrate that scAR is an accurate tool for protein denoising in CITE-seq.

### scAR accurately removes ambient mRNA in scRNAseq

Many methods have been developed for denoising or normalising scRNAseq data^6,7,10,11,20,33–36^. To demonstrate the advances of scAR, we benchmarked it with the most state-of-the-art approaches, including DCA^10^ and scVI^11^. We selected a public dataset^26^ that sequenced pooled human HEK293T and mouse NIH3T3. In this dataset, transcripts from the other organism were unambiguously ambient contamination, providing a ‘ground truth’ for the evaluation. By unsupervised transcriptome clustering, we identified 7590 HEK293T cells, 8006 NIH3T3 cells and 697 mixed droplets, which contained both HEK293 and NIH3T3 cells (supplementary Fig. 8a).

Interestingly, we noticed that the compositions of ambient transcripts (α in Fig. 1b) varied between different subpopulations of droplets (supplementary Fig. 8b-d and Methods). When taking different populations to calculate ambient frequencies, scAR outputted slightly different estimated contamination rates (supplementary Fig. 8e). Overall, scAR could precisely predict the cross-species ambient signals (i.e., the mouse transcripts in human cells or human transcripts in mouse cells). In comparison, the prediction for inter-species ambient signals (e.g., ambient mouse transcripts in mouse cells) depended on the inputting ambient frequencies. This suggests that a precise estimation of ambient frequencies is key to noise reduction. In addition, estimation of ambient frequencies by averaging cells yields similar results as by averaging cell-free droplets; therefore, scAR package uses the transcript frequencies in cells as a default setting to simplify the usage of scAR. However, we recommend calculating ambient frequencies using well-defined cell-free droplets to achieve the highest denoising performance.

We benchmarked these two modes of scAR along with scVI and DCA. Both human and mouse cells exhibited low exogenous contamination in the raw counts (Fig. 5a). After denoising, we observed that two modes of scAR removed all cross-species contamination (Fig. 5a-b and supplementary Fig. 9a). Nearly all mouse transcripts were identified and removed in HEK293T cells, and almost all human transcripts were also identified and removed in NIH3T3 cells. On the other hand, the mouse and human transcripts ratio was maintained at 1:1 in the multiplets after denoising.

**Fig. 5.**
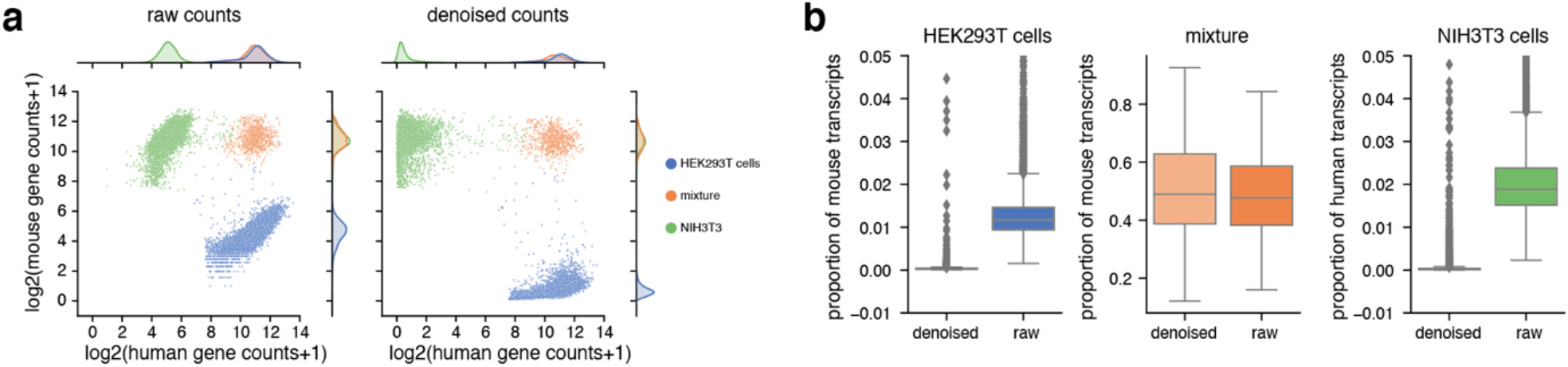
scAR eliminates ambient RNAs in scRNAseq. A public scRNAseq dataset of mixed human HEK293T and mouse NIH3T3 cells (1:1) was selected to demonstrate scAR’s ability in noise reduction in transcriptome data. **a**, Scatterplots show transcript composition before (left) and after (right) scAR denoising in three populations, HEK293T, NIH3T3 and multiplets. X- and y-axis show transcripts exclusively mapped to the human or mouse genome, respectively. **b**, Quantification of exogenous contamination before and after denoising in three populations. Y-axis represents per cell fraction of exogenous transcripts, i.e., mouse transcript rate in all HEK293T cells and human transcript rate in all NIH3T3 cells.

In contrast, DCA and scVI did not remove the cross-species contamination (supplementary Fig. 9b-c). The ambient signal hypothesis might partially explain this. Ambient signals were not random but ‘evenly’ distributed (the presence of ambient transcript follows a fixed probability per transcript) in each cell. Therefore, methods, like scVI and DCA, which specialised in reducing stochastic noise, might treat the ambient signals as the baseline ‘native’ signals.

### scAR is scalable and computationally efficient

Next, we benchmarked the resource efficiency by comparing the model size and runtime between scAR and similar methods. The model size (number of total parameters) is approximately linearly correlated to the number of features. Under default settings (e.g., default width and depth of neural networks), scAR has much fewer trainable parameters than totalVI, and scVI (~0.26x and ~0.54x, respectively), meaning scAR requires less memory resource and consumes less electric power. An exception is DCA, which has ~74% fewer parameters than scAR (Fig. 6a). However, training DCA is approximately ~55% slower than scAR under the same settings (e.g., sample size, epochs, and batch size) (Fig. 6b). It should be noted that DCA was trained on a 28-core CPU. GPU implementation of DCA can significantly shorten the training time. scAR is slightly faster than scVI, and it saves ~16% time in all test samples. For processing CITE-seq data, totalVI is faster than scAR as it simultaneously processes protein and mRNA data. In contrast, scAR needs to be run separately for protein and mRNA data (data not shown). Like DCA, scVI, and totalVI, scAR uses the minibatch functionality and can process a small number of cells at a time without scarifying accuracy. This makes it scalable to large-scale single-cell data even with limited GPU memory.

**Fig. 6.**
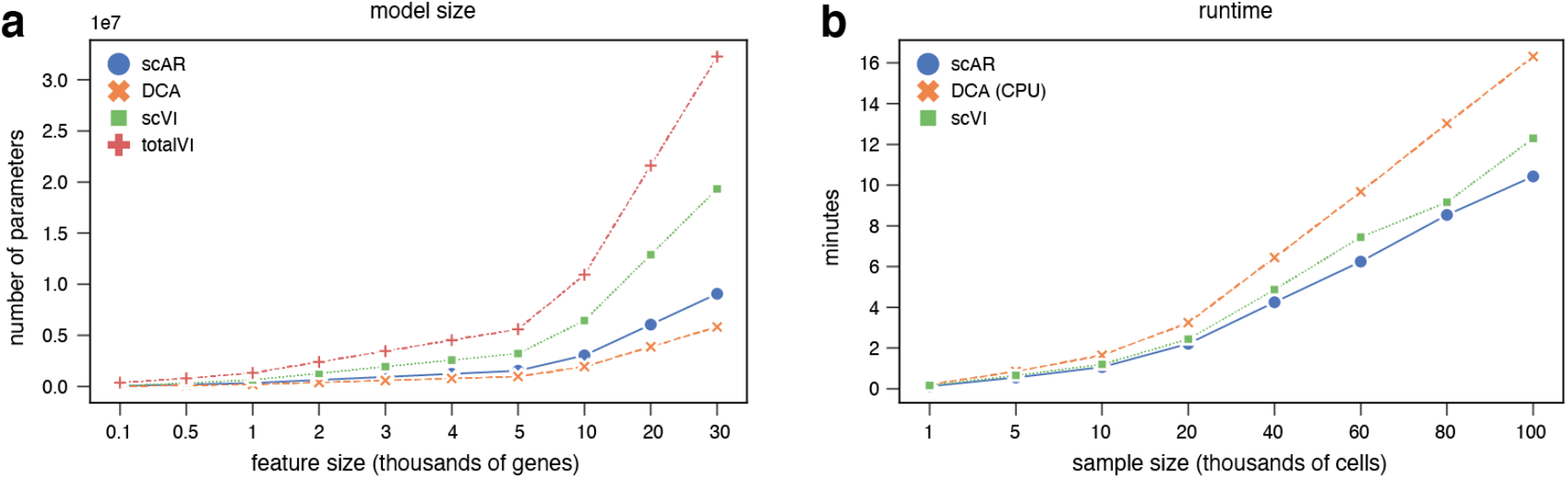
Benchmarking the efficiency of scAR. **a**, The lineplot shows the number of total trainable parameters under the default settings of each algorithm. **b**, The lineplot shows the runtime on subsampled cells of a 1.3 million single-cell dataset^26^. Experiments are run on the same machine and under the same parameter settings where possible.

## Discussion

Versatile single-cell omics technologies have expanded understanding of single-cell biology, and the development of new technologies is constantly pushing the boundary further. However, the noise In this context, we developed scAR to provide a reliable ‘one-for-all’ denoising solution for these technologies.

scAR can precisely infer the native signals for protein data in CITE-seq and mRNA data in scRNAseq. Recent approaches^11,12,20,27,33,37^ introduce deep learning technologies (such as AE and VAE) for these tasks and show great promise. These approaches generally design noise models based on data patterns (e.g., zero-inflation) and learn the model parameters through neural networks^11,13,33^. In scAR, we constrain the noise model under the ambient signal hypothesis and empirically estimate a set of parameters from cell-free droplets, this roughly reduces the outputs of VAE by half (supplementary Fig. 10) and focuses the VAE on learning the other latent variables, including biology-related native expression and the noise ratios. As a result, this hypothesis-driven modelling performs better at cleaning ambient signals than those that disregard the ambient signal hypothesis (Fig. 5 and supplementary Fig. 10). Moreover, it generalises scAR to fit a broader range of droplet-based single-cell datasets, independent of the data sparsity. However, it should also be noted that other methods, particularly DCA and scVI, has stronger strength in reducing stochastic noise and cellular variation than scAR because of their more flexible noise models (data not shown). Strictly speaking, scAR is distinct from these methods and can even be combined to achieve higher denoising performance. For example, one can consider scAR as a part of the pre-processing pipeline to remove ambient noise and subsequentially use other methods to reduce single-cell variance.

Another use case of scAR is the assignment of identity barcodes. Several single-cell omics technologies integrate exogenous unique barcodes into single-cell genome to label and/or perturb the host, such as single-cell CRISPR screens (e.g., CROP-seq, Perturb-seq and CRISP-seq)^1,2,28,29^ and cell indexing^14–16^. This class of single-cell technologies suffer heavily from the ambient noise of identity barcodes. Unlike CITE-seq and scRNAseq, the matrices of identity barcodes contain no signatures as protein or transcript data for cross-validation (e.g., denoising of a given marker can leverage information of other protein markers). Most of the current studies have assigned exogenous barcodes by hard filtering approaches^28,30^, which filter out cells with low depth and perform naïve assignment afterwards. This is not only inaccurate (Fig. 3) but also inefficient as it can discard as many as half of the cells despite their high quality of transcriptome readouts^38^. Other approaches such as MUSIC^38^ and scMAGeCK^39^ proposed to link transcriptome profile to sgRNA assignment. These methods are specific to single-cell CRISPR data (unlikely suitable for the cell indexing case). On the other, there is a risk of being misled by potentially dominant transcriptional states (e.g. cell cycle). They might also fail in other cases, for example, when several nodes of the same pathway are being interrogated or when the phenotypic effect of the perturbation is low (in the case of, e.g., low effective sgRNAs or wrong time points). In contrast, scAR uses only the information of identity barcodes (i.e., sgRNAs and cell tags) and evaluate the probability of ambient contamination for each barcode. This ensures a more accurate and unbiased assignment of identity barcodes. We tested scAR on single-cell CRISPR screening, but it should fit other technologies in this class as these technologies all use similar protocols to prepare, construct and sequence the barcodes (sgRNAs or identity barcodes).

Besides ambient contamination, other technical factors^8,9^ can also introduce background noise. We assume in scAR that ambient source is the most predominant artefact, and in turn, this hypothesis seems to be confirmed by the outstanding performance of scAR. However, further experimental validation may still be required. In addition, in CITE-seq technology, the non-specific binding of antibodies may bring in extra noise^4,13^. This is not modelled in scAR as we consider it too specific (dependent on the antibodies and experimental cell lines) to violate the scope of scAR’s generality. Moreover, identification of this noise may require dedicated well-designed experiments (e.g., spike-in^4^), as models can hardly distinguish between specific and non-specific binding without human intervention.

Finally, we observed different contamination levels in different datasets. scAR’s ability to estimate noise ratio may allow to evaluate batch effects and guide the experimental design, such as the protocols for cell fixation and washing. Furthermore, scAR has great potential in facilitating technology implementation and development in droplet-based single-cell omics. For example, single-cell analysis of accessible chromatin (scATAC-seq) also suffer from noise^40,41^, the concept of scAR might help design methods for the background correction. The second example might be scifi-RNA-seq^42^, which leverages cell indexing technologies to encapsulate and sequence multiple cells in a droplet to achieve ultra-high-throughput. One can imagine intense noise in this complex setting, where scAR may be beneficial for signal denoising.

## Methods

### The scAR model

scAR uses a hypothesis-driven probabilistic deep learning approach to infer the biological and technical variation in droplet-based single-cell omics experiments. The observed raw counts are modelled as a combination of biological signals and technical artefacts.

#### Modelling count data using Binomial regression

We take a generative approach to modelling the observed count matrix *X* ∈ ℕ_0_^*M*×*G*^, which denotes *M* cells and *G* features (i.e., genes, antibodies, sgRNAs or identity barcodes). A graphical model representation of this generative model is summarised in supplementary Fig. 11. For a given cell *m, x*_*m*_ represents a G-dimensional vector of observed expression data. We assume that *x*_*m*_ is drawn from a Multinomial model:

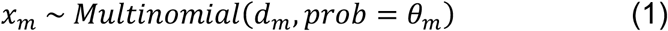

where *d*_*m*_ is the library size of cell *m* and *θ*_*m*_ is the feature frequency vector of size G. *d*_*m*_ is expected to be large enough to use a Poisson approximation for the multinomial. Therefore, for feature *g* in cell *m*, the observed count *x*_*mg*_ is drawn from a Binomial model:

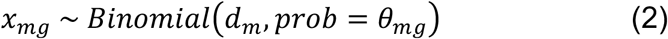

where *θ*_*mg*_ represents the probability of observing feature *g* in cell *m*. It is determined by two factors, the native expression *n*_*mg*_ and the ambient signals *a*_*mg*_, which can be modelled as,

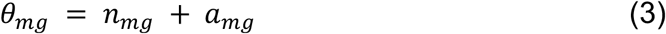

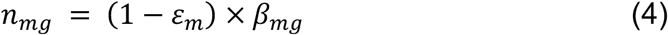

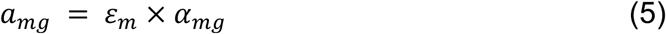

where *ε*_*m*_∈ [0,1] is a hidden variable, representing the fraction of total ambient counts. *β*_*mg*_ is another hidden variable, representing the feature frequency of the true native expression of feature *g* in cell *m. α*_*mg*_ represents the ambient frequency and according to ambient signal hypothesis, it is independent of cells, so we get,

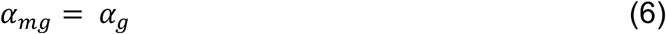

Notably, the cell-free droplets can be expressed as in equation (2), with the native component being zero and noise ratio being 1, so,

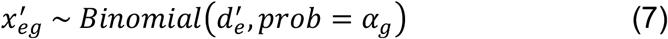

Where, 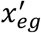 and 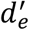 represent counts for feature *g* and library size in cell-free droplet *e*, respectively. According to the law of large numbers, we can approximate *α*_*g*_ by averaging feature counts in cell-free droplets,

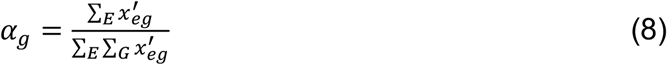

Put all together, we have,

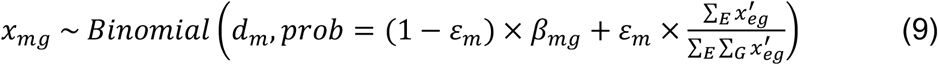

where only *ε*_*m*_ and *β*_*mg*_ are unknown parameters that need to be estimated. According to Bayes’ theorem, we get the posterior probability,

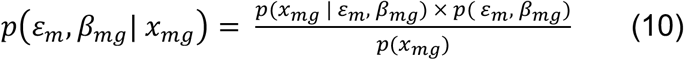

Since the prior probability *p*(*ε*_*m*_, *β*_*mg*_) and likelihood *p*(*x*_*mg*_ | *ε*_*m*_, *β*_*mg*_) are both generally intractable, we implement a variational inference^23,43^ approach to estimate *ε*_*m*_ and *β*_*mg*_, as described in the following section. To ensure flexibility, we also provide implementations without the Poisson approximation to allow users to choose and test.

#### Variational inference for scAR

We apply variational autoencoder strategy to estimate the hidden variable *ε*_*m*_ and *β*_*mg*_ mentioned above. The architecture of the VAE is demonstrated in supplementary Fig. 12. VAEs introduce an additional latent variable z in the bottleneck layer so the marginal log-likelihood of observation *x*_*m*_ can then be written as,

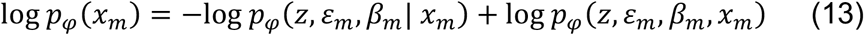

where *φ* represents the parameter space, i.e., model weights. *ε*_*m*_ and *β*_*m*_ are calculated by a deterministic neural network (i.e., the decoder),

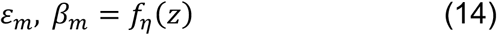

where, *f* represents the decoder and *η* ⊂ *φ* represents the trainable weights of *f*. This means,

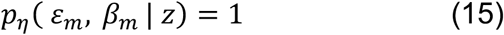

Therefore, we can integrate out *ε*_*m*_ and *β*_*m*_ and re-write the equation (13) as

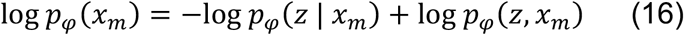

We construct variational posterior *q*(*φ* | *ω*) to approximate the posterior *p*(*φ* | *x*_*m*_). Therefore, we have,

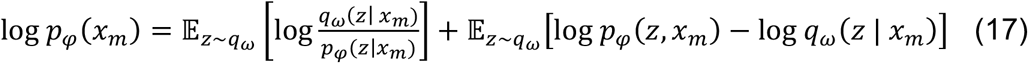

where the first term on the right side is the Kullback-Leibler divergence^44,45^ between distributions *q* and *p*, reflecting the difference between parameter distributions. It is non-negative, so we can get the evidence lower bound (ELBO) as follows,

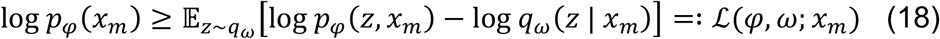

Increasing the ELBO will approximate the distribution *q* to *p* thereby ensuring the learnt variables are as close as the expectation. Therefore, the ELBO is generally used as the objective function to fit the VAE. We can further transformation equation (18) into,

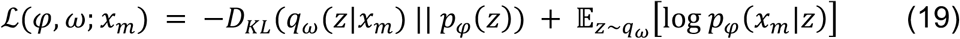

The negative of ELBO is used as loss function to simultaneously optimize model weights and hidden variables in scAR. In case of M cells, the loss function is then written,

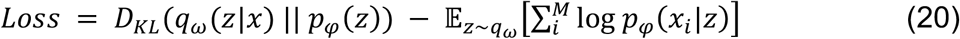

Minimizing the loss function requires a tradeoff between the KL divergence and expected negative log-likelihood term. On the one hand, the KL divergence between *q*_*ω*_ (*z*|*x*) and *p*_*φ*_(*z*) should be kept small, preventing the variational posterior from being too different to the prior. On the other, the variational posterior parameters should maximize the log likelihood log *p* _*φ*_(*x*|*z*), ensuring a small reconstruction error of scAR. We use the reparameterization trick to calculate the gradients with respect to *φ* and *ω* for KL term^45^. According to equations (14) and (15), we have,

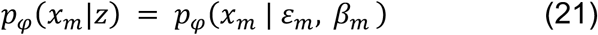

Since we assume *x*_*m*_ is drawn from Binomial distribution with latent parameters *ε*_*m*_, *β*_*m*_ (see equation (9)), *p*_*φ*_(*x*_*m*_ | *ε*_*m*_, *β*_*m*_) also has a closed-form expression, thus the gradient descents of negative log-likelihood term in equation (20) are easy to calculate. Together, we use the gradients of the loss function to update the parameters *φ* and *ω* to determine the hidden variables noise ratio *ε* and expected native frequencies *β*.

#### Bayesian inference and assignment of identity barcode

We infer the expected native signals 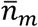 and ambient signals *ā*_*m*_ in cell *m* using the following equations,

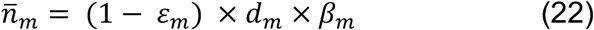

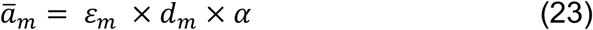

where 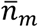 is used as the denoised counts.

Furthermore for the assignment of sgRNAs in CROP-seq, we also use and provide Bayesian factor as a metric to compare two hypotheses: the observed counts consist of both native and ambient sgRNAs (*H*_1_) vs the observed counts contain only ambient sgRNAs (*H*_2_). For a given sgRNA *g* in cell *m*, this can be mathematically expressed as follows,

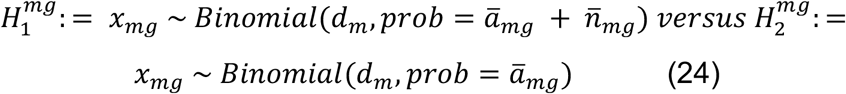

The Bayesian factor is then given by,

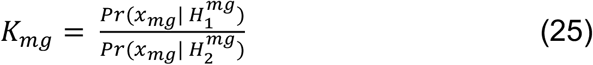

The numerator and denominator represent the probability that *x*_*mg*_ is produced under assumption of 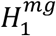 and 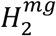, and we approximate them using the cumulative distribution function (stats.binom.cdf) and probability mass function (stats.binom.pmf), respectively. High *K*_*mg*_ (>=3) favours the first hypothesis, meaning the sgRNA *g* contains native signal. In the case of multiple high *K*, we assign the sgRNA of highest *K* to the cell.

#### Model optimisation for scAR

To identify a universal set of hyperparameters as the default setting for scAR, we perform grid search on two types of synthetic datasets (see supplementary note I), which simulate CROP-seq data type and CITE-seq/scRNAseq data type respectively. To limit the number of parameters, we fix several less important parameters. For example, the training epochs are fixed at 800. Additionally, we use the Adam optimizer^46^ with exponential decay to schedule the learning rate but the decay rate is fixed at 0.97 every 5 epochs. The hyperparameters which are optimised include the number of nodes of neural networks, the dimension of latent space, the dropout probability of neurons, the initial learning rate, and the KL divergence weight. As a result, we tested 6912 combinations of parameters for each dataset (supplementary Fig. 13) and identified the most appropriate set listed as follows: units of 1st layer: 150; units of 2nd layer: 100, dimension of latent space: 15; initial learning rate: 0.001, dropout probability: 0; KLD weight: 1e-5. All experiments were performed using these optimised parameters unless otherwise specified.

### CROP-seq experiment

The CROP-seq library was cloned into a modified pLKO-TET-ON plasmid in a pooled format by Golden Gate. The cloning reaction product was used to transform Endura electrocompetent cells, which were expanded in LB medium overnight (OD600 = 0.8) and plasmid DNA was harvested using Genopure plasmid maxi kit (Roche). We produced lentiviral particles and transduced MCF7-dCas9-KRAB cells (MOI = 0.3) with the CROP-seq library. The cells were selected with 2μg/ml puromycin (Invitrogen) and they harvested at defined time points by FACS (mCherry-positive cells). The single-cell suspensions were fixed in 90% methanol in DPBS (v/v) and stored at −80 °C prior to rehydration and further processing. The rehydration buffer was supplemented with 1% Bovine serum albumin and 0.5 U/ul RNase inhibitor (Sigma, P/N 3335399001). All samples were processed using Chromium Next GEM single-cell 3’ reagents kit (10x Genomics) according to the manufacturer’s protocol and the libraries were sequenced in an Illumina HiSeq 2500.

### Pooled CRISPR screen

MCF7-CRISPRi cells were transduced with independent lentiviral pools (MOI = 0.3) of the CROP-seq library. To guarantee a correct representation of all sgRNAs in the cell population we transduced ≈1000 cells per plasmid. The cells were selected using 2μg/ml puromycin (Invitrogen) at 24 hours post-transduction, after which they were expanded and harvested at indicated time points. We extracted gDNA from the cells using DNeasy kit (Qiagen) and prepared libraries for next generations sequencing.

### Analysis of CROP-seq data

Single-cell sequencing data were processed using Cell Ranger (version 3.1.0, 10x Genomics) and sgRNA count matrices were generated using KITE (https://github.com/pachterlab/kite). Human genome assembly (Ensembl GRCh38 release-98) was used as the reference to map mRNA reads. The Scanpy package^47^ was used to perform quality control, cell filtering, gene filtering and differential expression analysis. We used two normalisation approaches to examine the knockdown effect. The first one as shown in Fig. 2g-i is a library size normalisation. Sequencing depth per cell was normalised to 1.0xe^5^ counts and t-test was performed on the normalised counts across cell groups using the scipy.stats.ttest_ind function. The second one (supplementary Fig. 3a) is a Z-normalisation as reported in our previous publication^3^. For each gene, we subtracted the mean value of CTL group then divided by the standard deviation of the CTL group.

### Analysis of CITE-seq data

The cellranger outputs of PBMCs5k^26^ dataset were downloaded from 10x genomics. Cells with extreme counts (<1500 counts or >15000 counts) were discarded. Stressed cells with high presence of mitochondrial genes (>=0.2) were also discarded. The cell clustering was performed using Scanpy and annotated based on expression of a panel of marker genes.

Correlation of RNA-protein pairs. Spearman’s correlation was performed between RNA and protein counts using scipy.stats.spearmanr function. Control antibodies were removed for this correlation analysis. CD45RA and CD45RO, which are encoded by an identical gene PTPRC were also removed due to the difficulty of identifying isoform transcripts. In addition, several markers (CD15, CD34, CD80, CD137, CD274, CD278, PD-1) were removed due to extremely low counts of either protein or corresponding RNAs.

### Species-mixing experiment

The cellranger outputs of species-mixing dataset^26^ were downloaded from 10x genomics. Scanpy was used to perform quality control, gene filtering, cell filtering and species identification. In brief, we first took the ‘filtered_feature_bc_matrix’ from cellranger output and further filtered out genes with extreme counts (<200 or >6000 in total) and cells with low gene counts (<200). We then performed library size normalization, log transformation, clustering and UMAP. By checking the differently expressed genes, we identified 7590 HEK293T cells, 8005 NIH3T3 cells as well as 697 multiplets mixed with both HEK293T and NIH3T3.

Examination of droplets. To identify the best representation of ambient signals, we examined subpopulations of droplets in the unfiltered matrix – namely, ‘raw_feature_bc_matrix’. All droplets were ranked by their total UMI counts and split into four subgroups through kneeplot: 1) droplets in ‘filtered_feature_bc_matrix’ were marked as cells, 2) droplets with high counts (>40) were marked as ‘droplet I’, 3) droplets with intermediate counts (>12 and <=40) were marked as ‘droplet II’, 4) droplets with low counts (<=12 and >0) were marked as ‘cell-free droplets’. We took the total gene frequencies in each subpopulation as the ambient frequencies and run scAR to compare the estimated noise ratio.

### Benchmarking efficiency

For benchmarking the model size, we generated synthetic CITEseq data (supplementary note I), run the models for 4 epochs, and counted the trainable parameters. For benchmarking the runtime, we downloaded a 1.3 million single cell dataset from 10x genomics, selected top 5000 highly variable genes, randomly sampled subpopulations of various sample size, and run the models with identical settings. We turned off the early stopping and fixed at 100 epochs with batch size of 128. We took 0.99 of the total samples as training set. Default values were used for other parameters. We run the benchmarking experiments using one 28-core Intel E5-2690 CPU (500 GB RAM), and one NVIDIA Tesla P100 GPU (16 GB RAM). scAR, totalVI, and scVI all were run on the GPU, and DCA were run using all 28-core CPU.

## Data availability

The CROP-seq data discussed in this manuscript have been deposited to the Sequence Read Archive and will be accessible through BioProject accession number: PRJNA794328. All other datasets are public. The CITE-seq datasets (PBMCs5k) and HEK293T and NIH3T3 pooled scRNAseq (20k_hgmm dataset) were downloaded from 10x genomics datasets. Other datasets were downloaded from Sequence Read Archive.

## Code availability

The scAR package is available at GitHub (https://github.com/Novartis/scAR). The full codes to reproduce the results in this manuscript will be available at GitHub upon publication (https://github.com/CaibinSh/scAR-reproducibility).

## Acknowledgements

We thank Joel Wagner, Joshua Korn, Viveksagar Krishnamurthy Radhakrishna, and Jeanne Whalen for inspiring discussions and testing scAR, Mathias Eder and Esther Uijttewaal for additional technical support.

## Author contributions

C.S. and A.d.W. conceived and designed the study. C.S., G.L. and A.d.W. designed the statistical model and performed analysis. R.L. and G.G.G. designed CROP-seq and bulk sequencing experiments and R.L. conducted the experiments. A.W. and R.C. performed scRNAseq and bulk sequencing and S.S. performed pre-processing of CROP-seq data. S.D., A.K., E.D., G.G.G, G.R. and A.d.W. supervised the study. C.S. wrote the original draft. C.S., G.L., R.L., S.D., E.D., G.G.G, G.R. and A.d.W. reviewed and edited the draft.

## Supplementary Table

**Supplementary Table 1.**
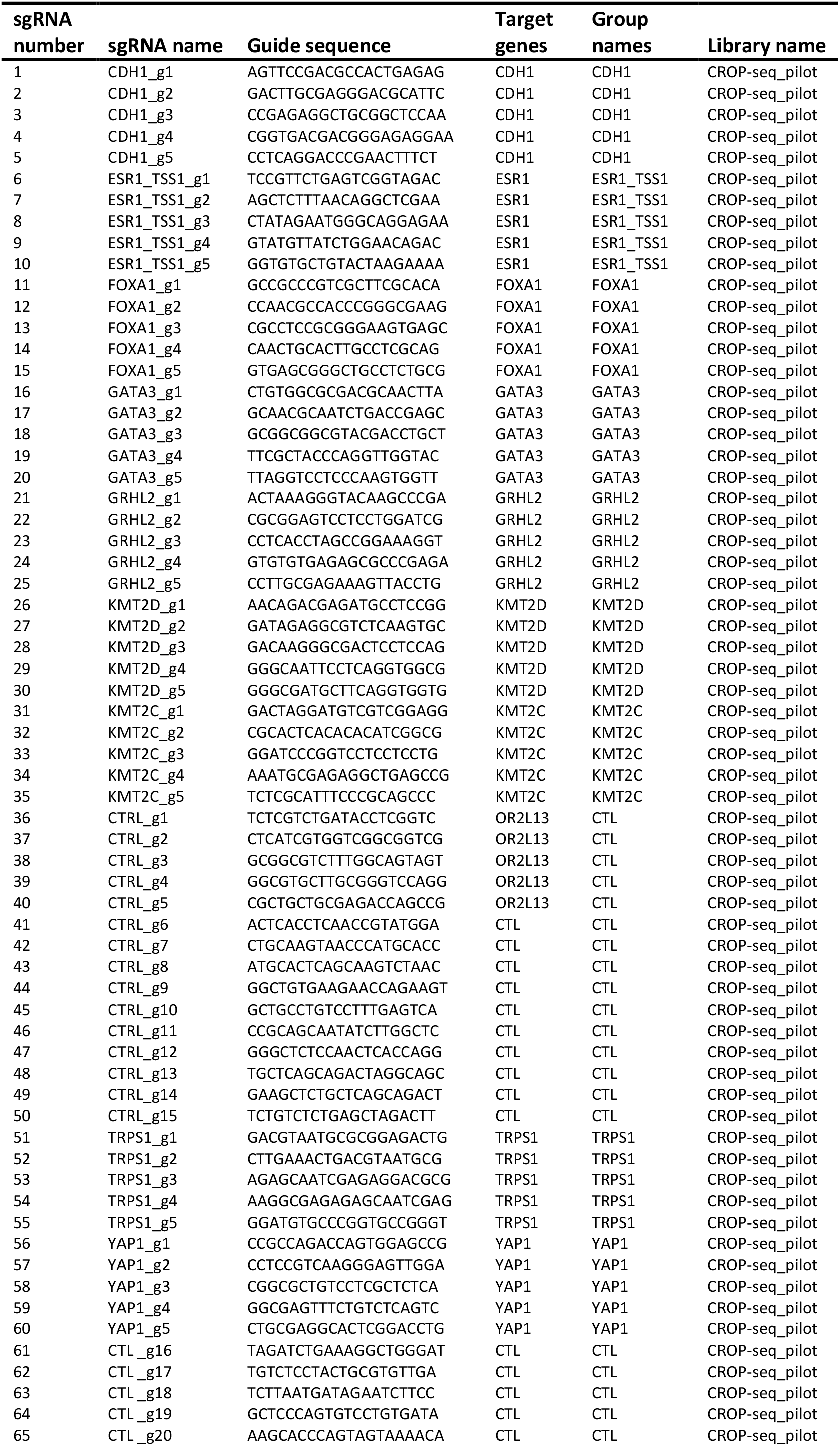
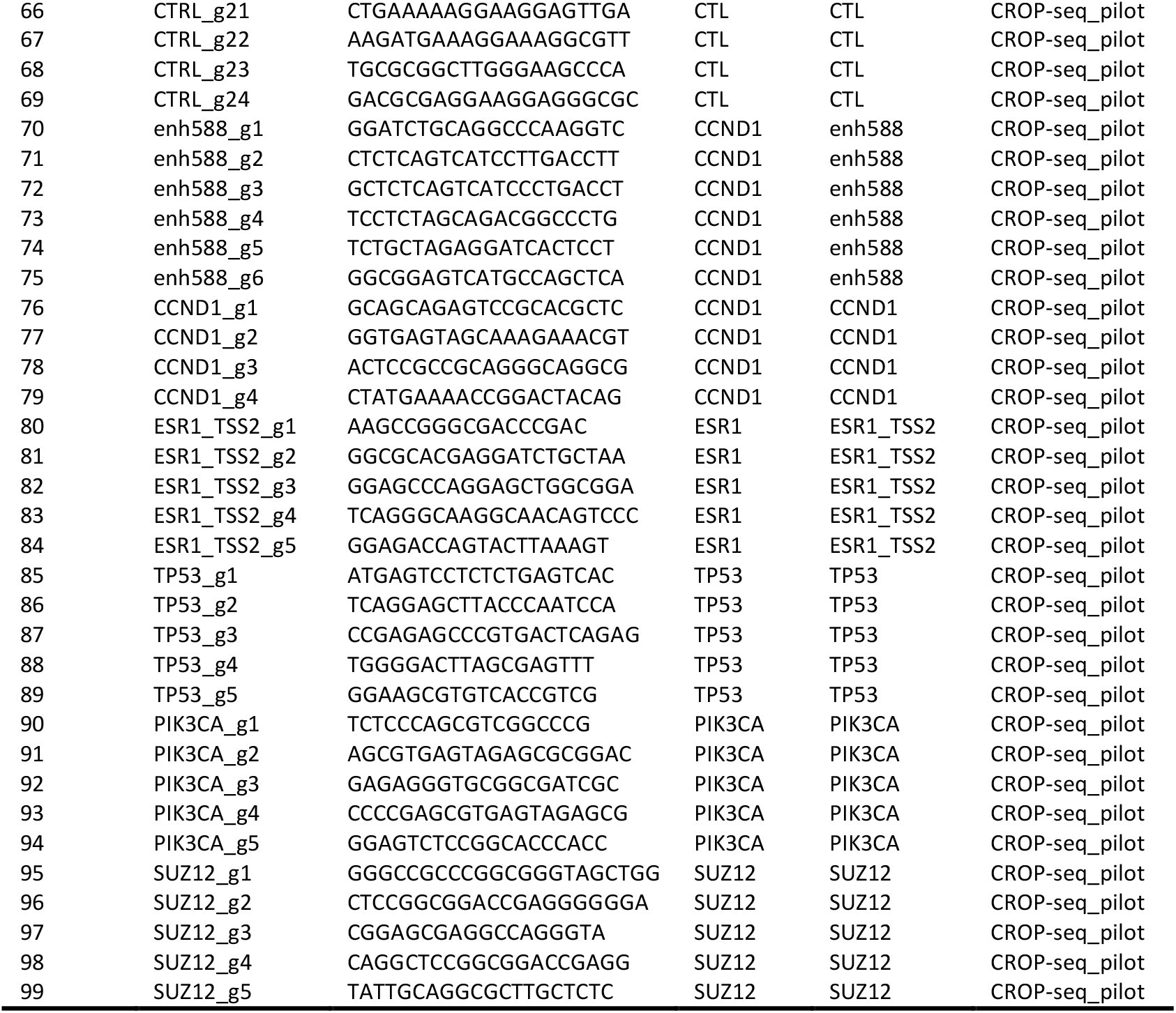
CROP-seq libraries.

## Supplementary Figures

**Supplementary Fig. 1.**
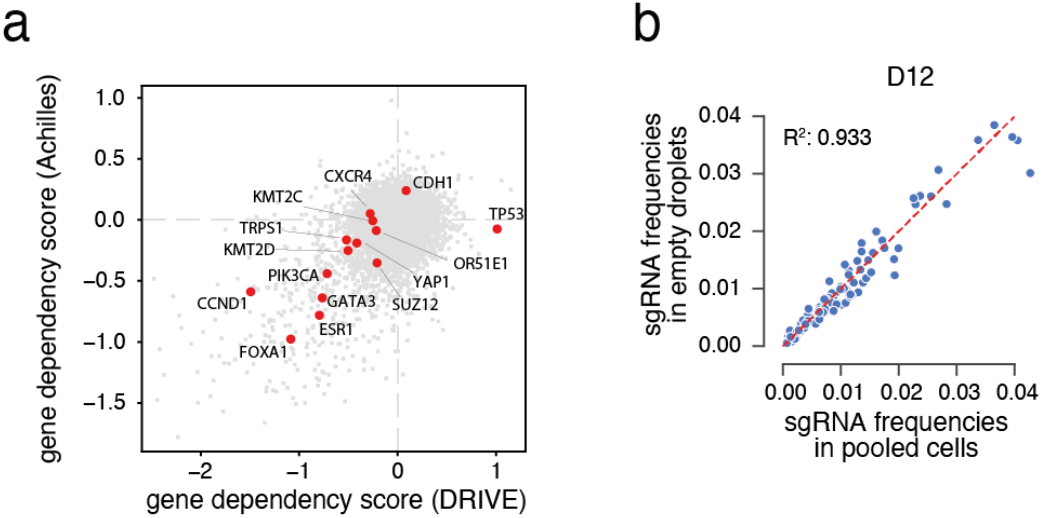
Validation of CROP-seq libraries. **a**, The gene dependency scores of most target genes show negative effects on the cell growth of MCF7. One exception is the tumour repressor TP53, down-regulation of which promotes tumour growth. Data are originally from two pooled screening projects (DRIVE and Achilles) and obtained from https://depmap.org/. **b**, Scatterplot of sgRNA frequencies between two independent experiments. The X-axis represents sgRNA frequencies in bulk DNA sequencing, Y-axis represents sgRNA frequencies obtained by averaging RNA counts in cell-free droplets from 10x scRNAseq. Each dot represents an sgRNA, the red dashed lines represent y=x, and coefficients of determination (*R*^2^ scores) is shown. D12 sample was shown

**Supplementary Fig. 2.**
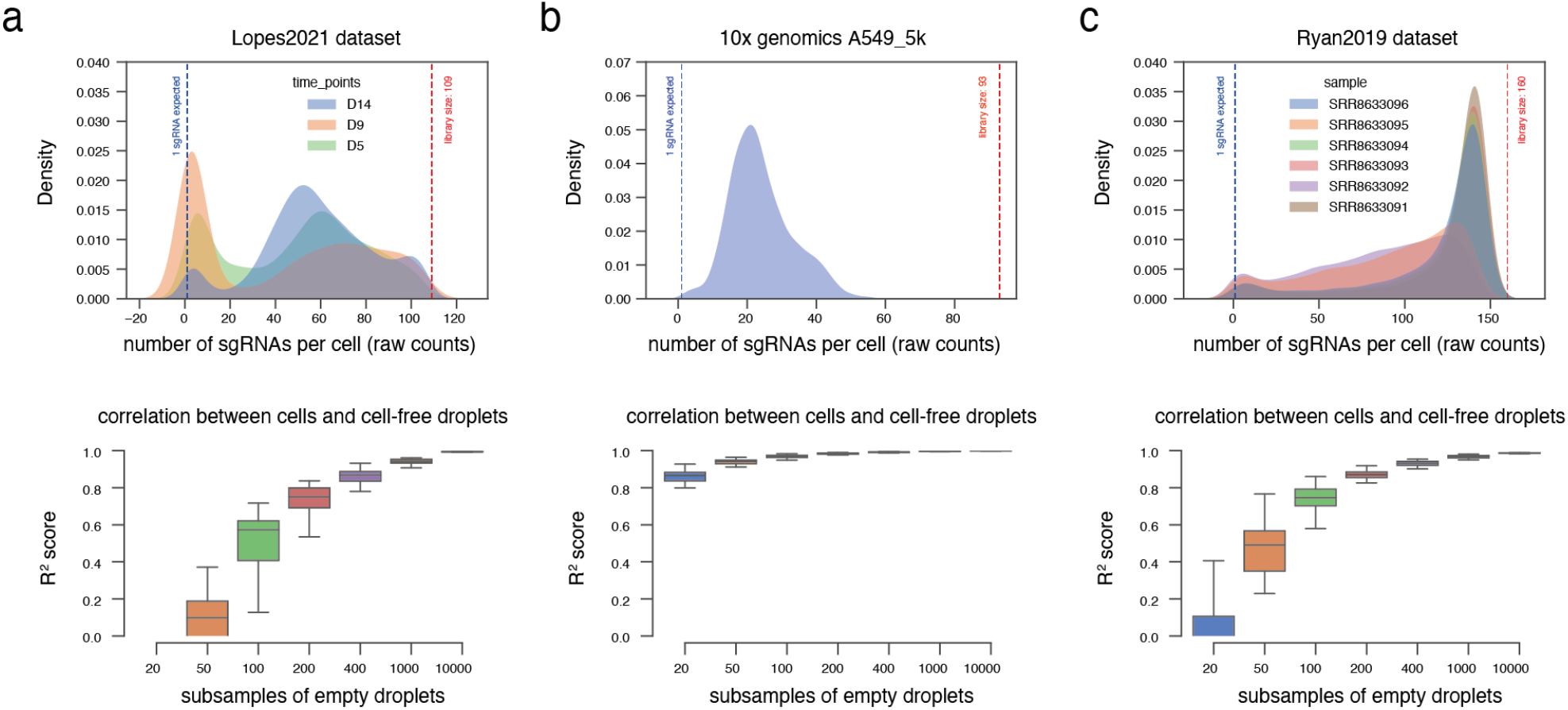
supplementary to Fig. 2, ambient noise widely exists in single-cell omics datasets. Ambient contamination is observed in several public datasets and the background noise is highly correlated with endogenous signals in cells. **a**, The Lopes2021 dataset^1^. **b**, The A549_5k dataset from 10x genomics^2^. **c**, The Ryan2019 dataset^3^.

**Supplementary Fig. 3.**
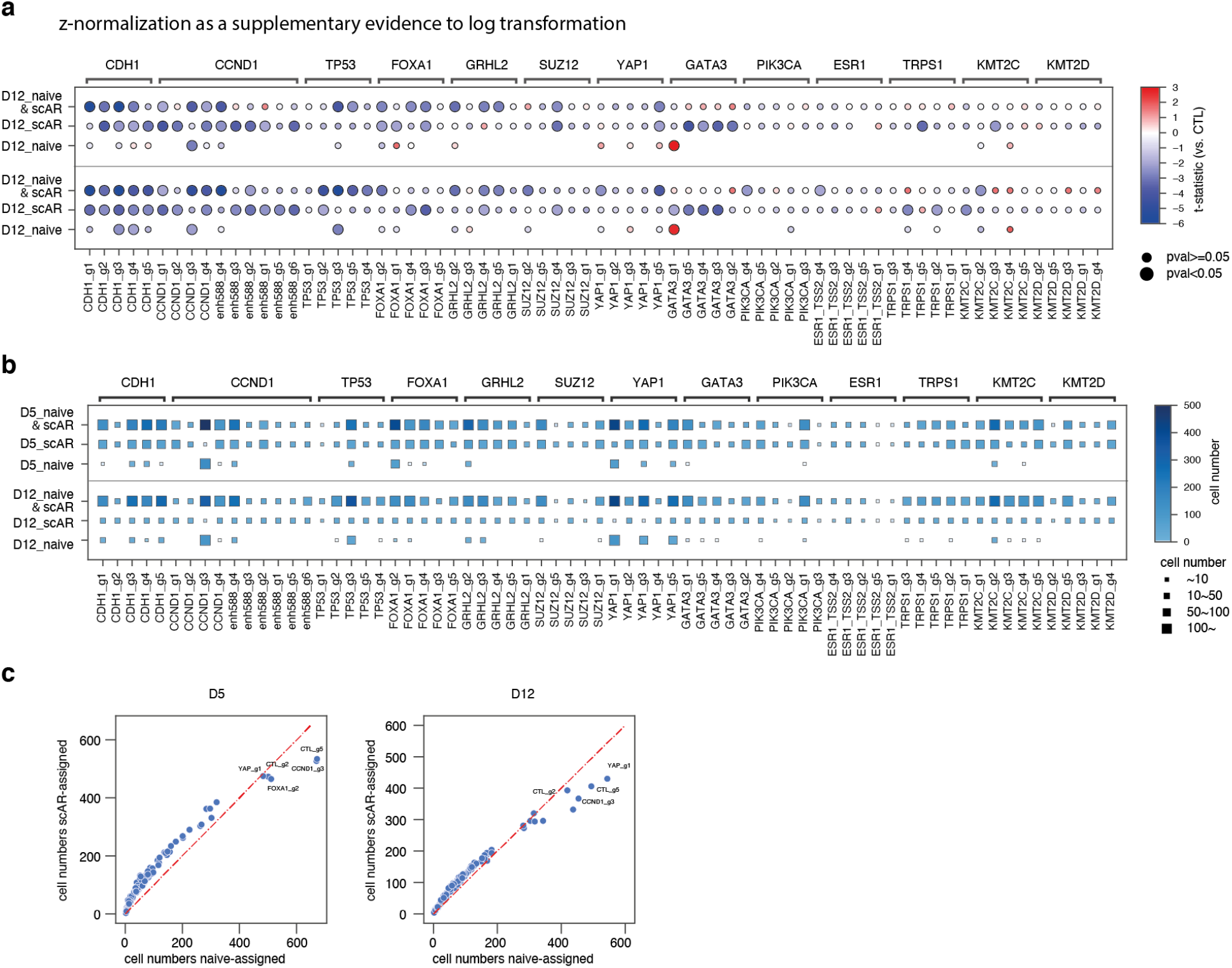
Supplementary to Fig. 3. Evaluation of scAR on CROP-seq. **a**, Similar to **Fig. 3g**, the dotplot shows the overall comparison between two assignment approaches on t-statics of z-normalized expression. The X-axis represents guide groups. Y-axis represents subgroups of cells as exemplified in **Fig. 3e** and **Fig. 3f**, separated by two time points and assignment approaches. Target genes are shown on the top. Their expression (log-transformed) in each group is compared with that in the CTL group and resulting t-statistics are shown by the dot colour. Blue colour indicates down-regulation, and red indicates up-regulation. CTL group is centred at zero. The bimodal sizes of circles represent the p-values from the t-test (the bigger means p<0.05, the smaller means p>=0.05). The asterisk (*) highlights the guide groups where scAR significantly improves the accuracy and hashes (#) mark the groups where scAR underperforms naïve assignment. **b**, The comparison between two assignment approaches on cell number after assignment. Sizes of squares represent the cell numbers of each assignment. **c**, The overall comparisons of cell numbers. Each dot represents an sgRNA.

**Supplementary Fig. 4.**
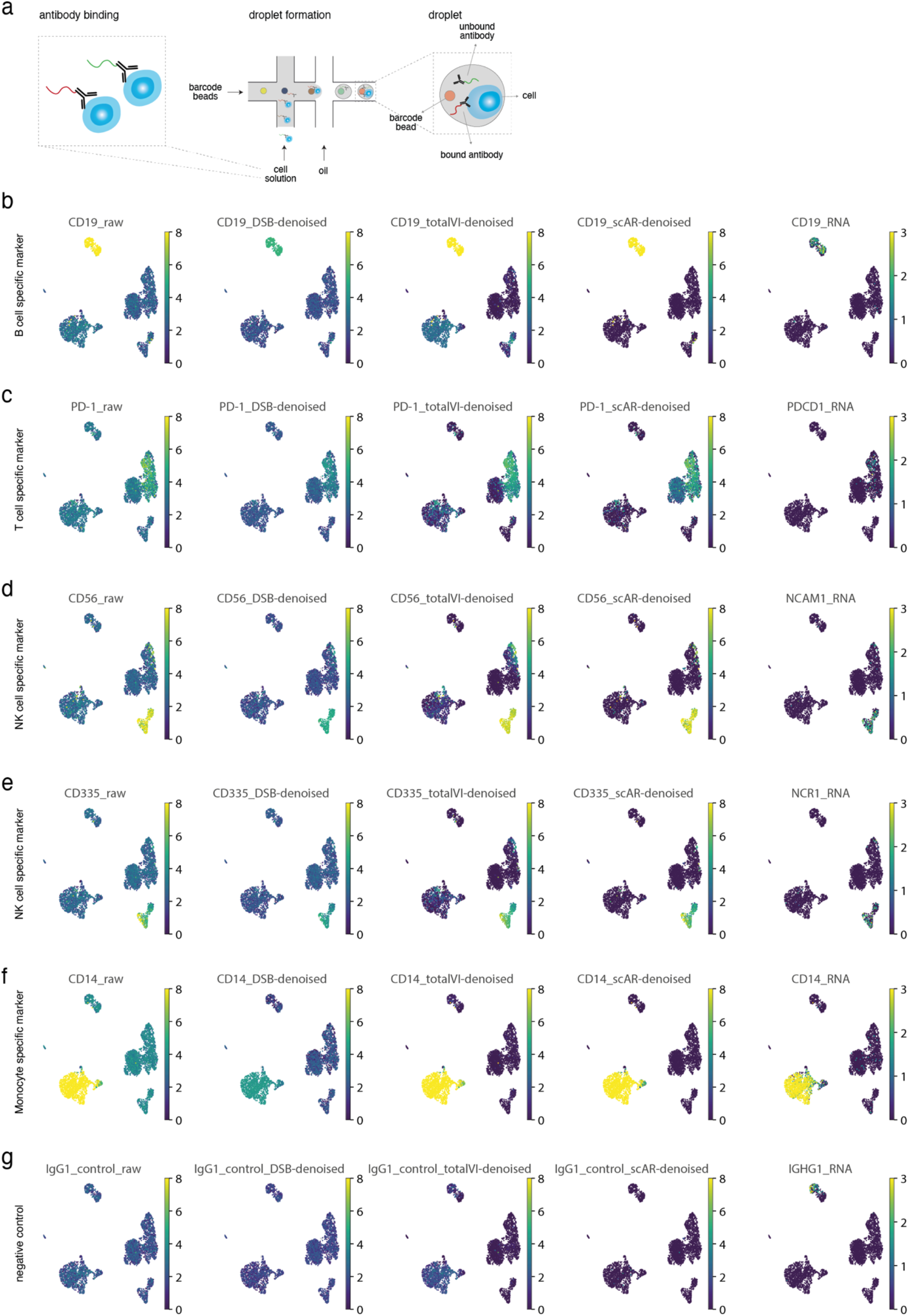
scAR reduces the non-specific ambient antibodies. **a**, The illustration of CITE-seq experiment and the potential source of background noise. **b-g**, UMAPs visualize raw protein counts, DSB-normalised, totalVI-denoised, and scAR-denoised protein counts, and the corresponding RNA counts of cell-specific markers, including a B cell-specific marker CD19 (**b**), a T cell-specific marker PD-1 (**c**), two NK cell-specific markers CD56 (**d**) and CD335 (**e**), and a monocyte-specific marker CD14(**f**), as well as a negative control IgG1 (**g**).

**Supplementary Fig. 5.**
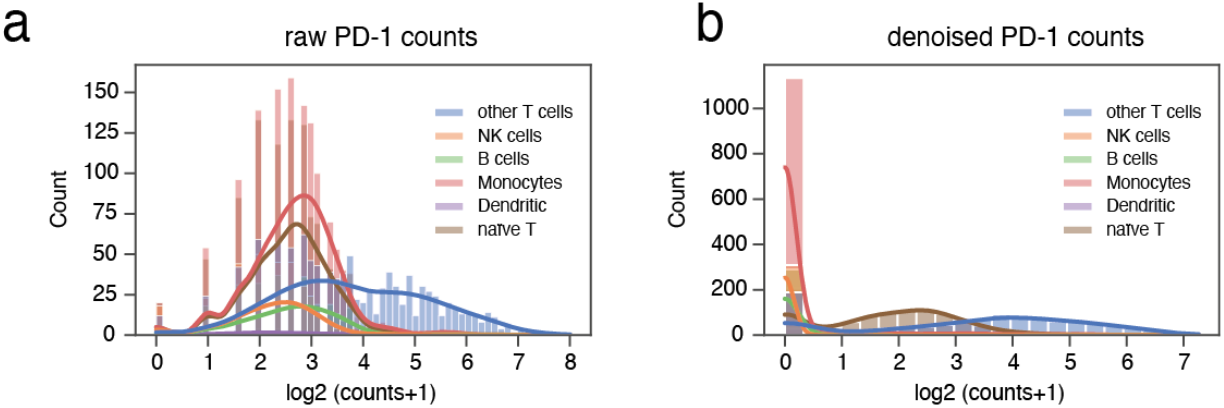
scAR removed the non-specific ambient PD-1 but preserved specific PD-1 even when they had similar counts. **a**, The distributions of raw PD-1 counts were similar/highly overlapped between cell types. **b**, scAR preserved true PD-1 signal in naïve T cells but removed false PD-1 signal in other cell types.

**Supplementary Fig. 6.**
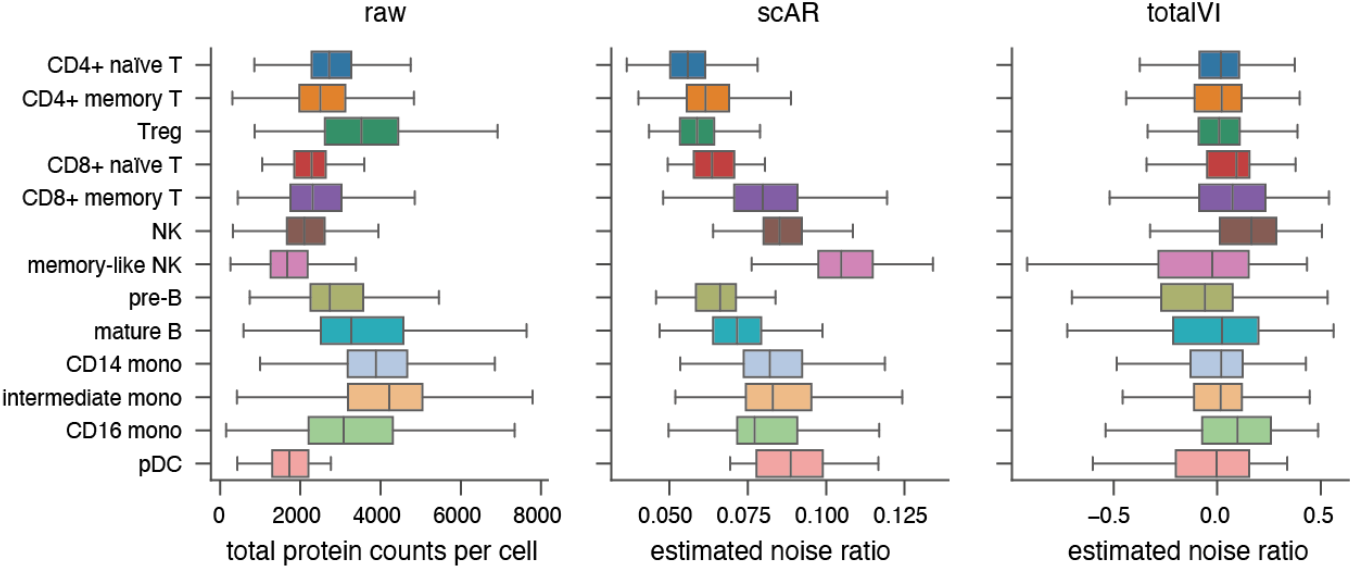
scAR removed <10% ambient counts across cell types and showed high consistency across cells. **a**, total raw counts of antibodies in cell subtypes. **b**, scAR-estimated noise ratio in cell subtypes. **c**, totalVI-estimated noise ratio in cell subtypes.

**Supplementary Fig. 7.**
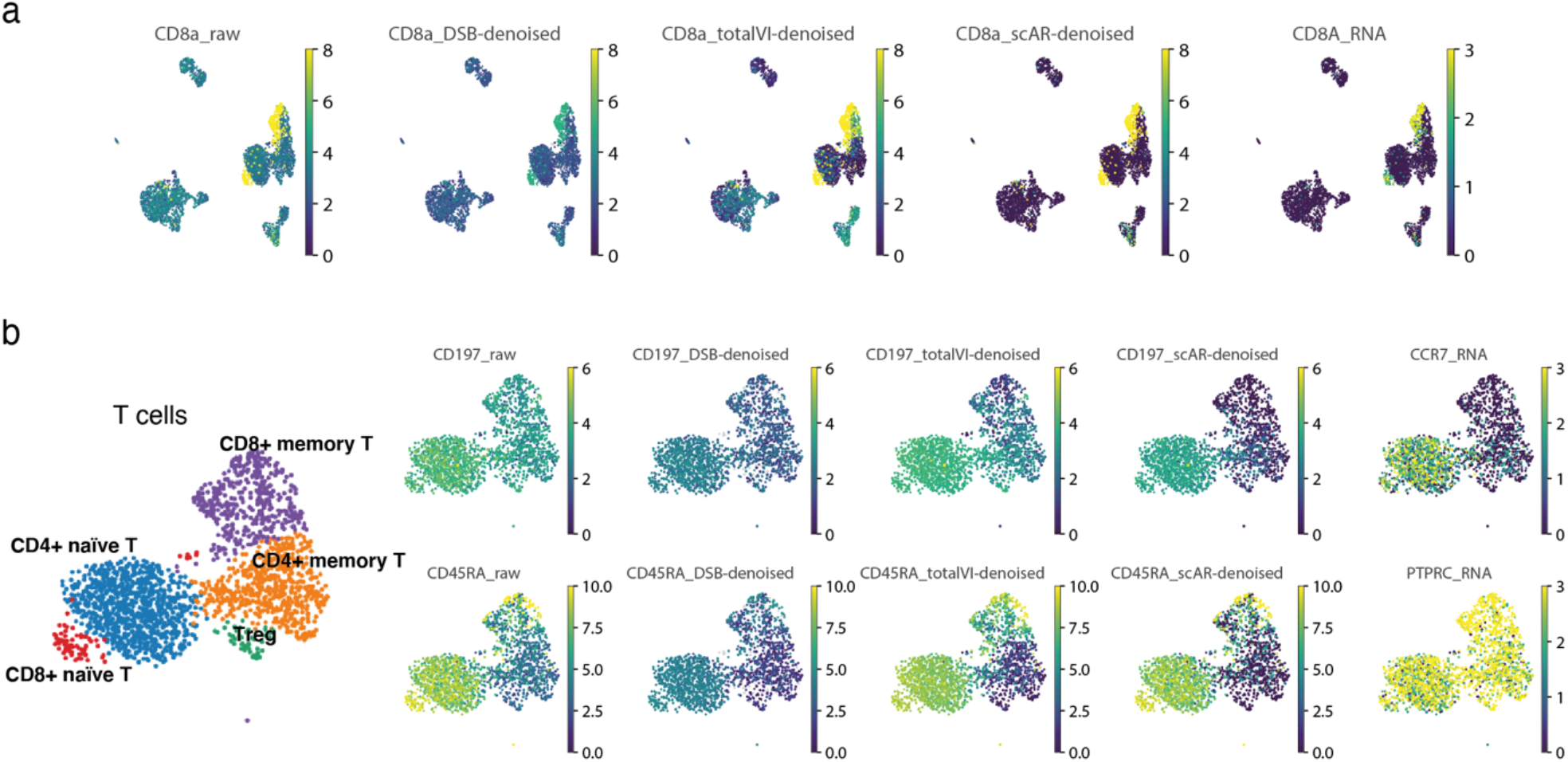
scAR denoised proteins show better separation of cell subtypes. **a**, scAR-denoised CD8a was exclusively expressed in CD8+ T cells. **b**, scAR-denoised CD197 and CD45RA separated better memory and naïve T cells.

**Supplementary Fig. 8.**
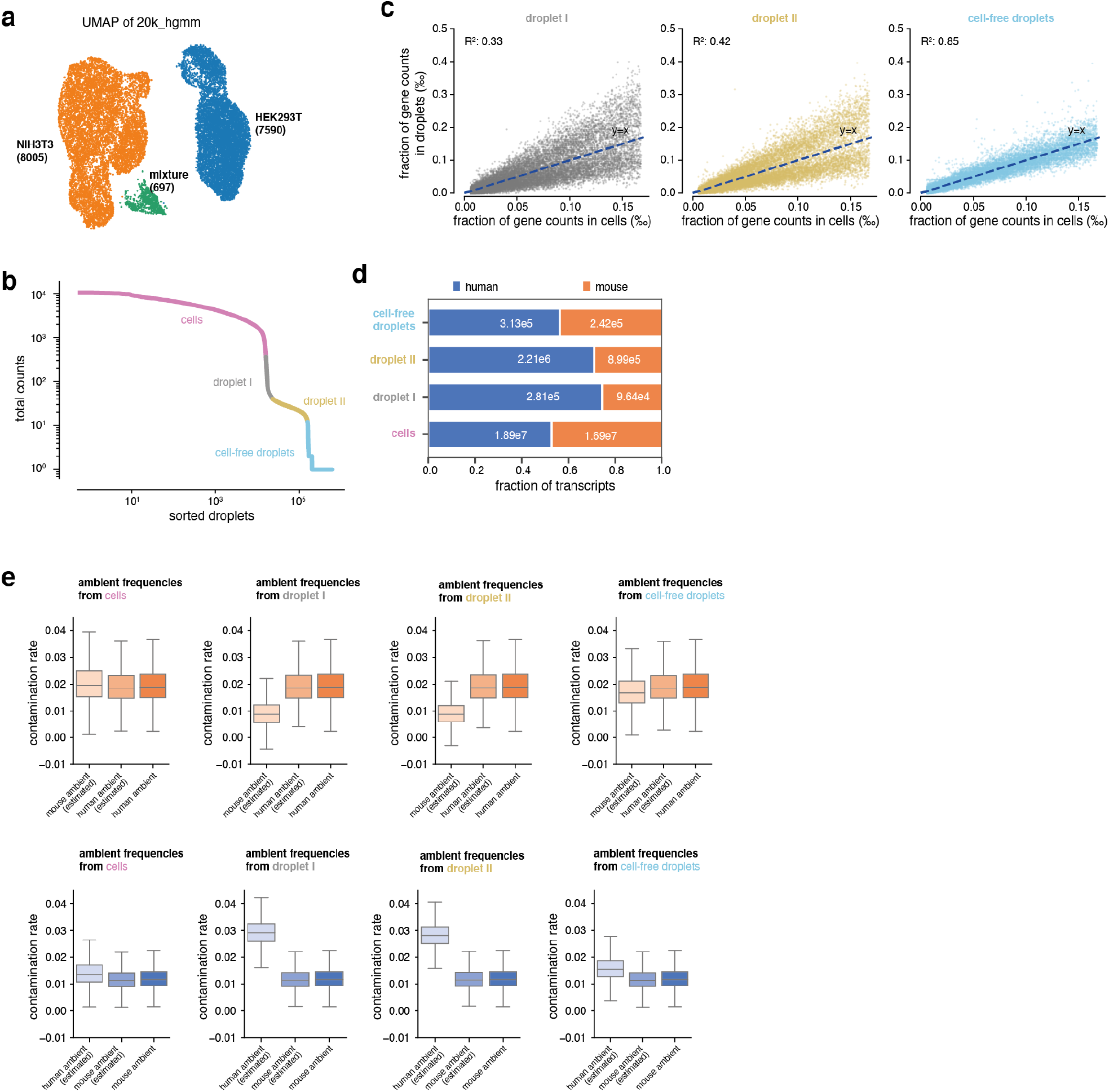
Supplementary to Fig. 5. A public scRNAseq dataset of mixed human HEK293T and mouse NIH3T3 cells (1:1) was selected to demonstrate scAR’s ability in noise reduction in transcriptome data. **a**, UMAP shows three populations of cell-containing droplets, HEK293T, NIH3T3 and multiplets. **b**, The kneeplot shows subpopulations of droplets. **c**, Correlation of gene frequencies between subpopulations of droplets and cell-containing droplets. **d**, Fraction of transcripts in subpopulations of droplets. **e**, Boxplots show the percentage of ambient signal in NIH3T3 cells (the upper panel) and HEK293T cells (the bottom panel). Columns separate the different inputted ambient frequencies which were calculated using different subpopulations of droplets. The Y-axis represents contamination ratio, the X-axis represent scAR-estimated inter-species, scAR-estimated cross-species, and true cross-species contamination, respectively. Given that the global ratio of human and mouse transcripts is ~1.11 in cells (**d**), it is reasonable to expect a similar inter-species and cross-species contamination ratios. However, droplet I and II lead to too high estimation of human-sourced contamination, as much as ~3x of mouse source. This may be explained by the over-representation of human transcripts in droplet I and II (**d**). The higher human transcripts in ambient frequencies as input, the more counts to be identified as background noise by scAR. The best estimate of ambient frequencies should be drawn from the population of cell-free droplets, as the estimated noise ratios are in a reasonable range in both cell lines – ambient signals from human sources are slightly stronger than mouse sources in both cell lines. These observations also suggest that compositions in these droplets are clearly different, e.g., droplet I and II may contain more human cell debris.

**Supplementary Fig. 9.**
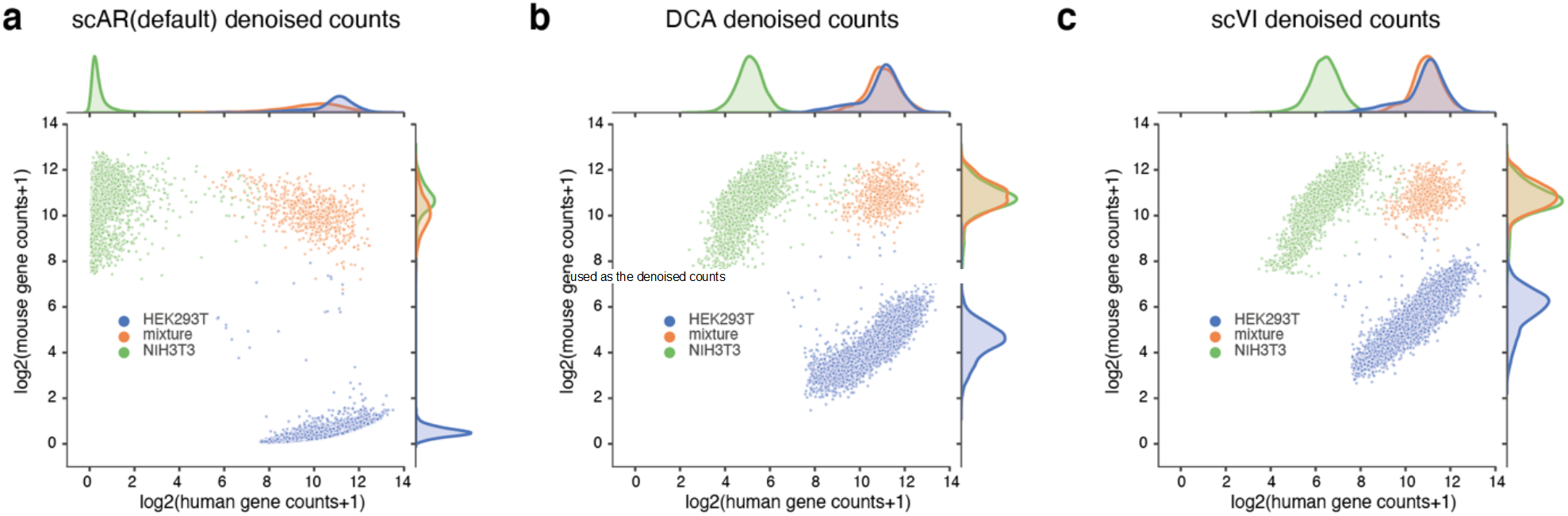
Supplementary to Fig. 5. Benchmarking of mRNA count denoising methods. **a**, scAR denoised mRNA counts under the default setting – ambient frequencies were calculated using cells. **b**, DCA denoised mRNA counts under its default setting (epochs and batch size were aligned to scAR at epochs = 800, batch_size = 64). **c**, scVI denoised mRNA counts under its default setting (epochs and batch size were aligned to scAR at epochs = 800, batch_size = 64).

**Supplementary Fig. 10.**
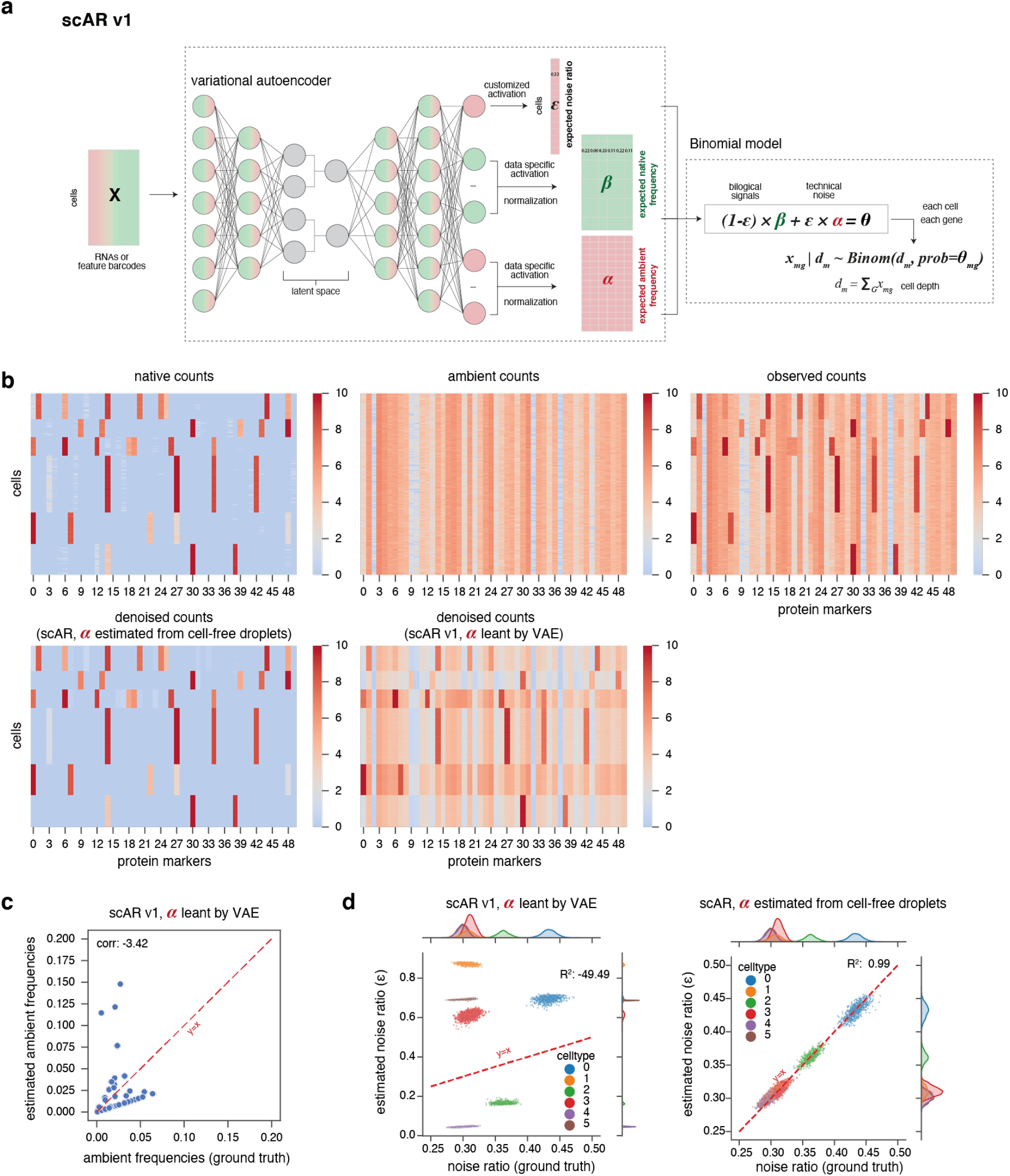
The performance of two versions of scAR. We compared two versions of scAR to demonstrate the necessary of using cell-free droplets. **a**, scAR v1 is an early version of scAR in which we fully relied on VAE to learn the ambient frequencies. **b**, Heatmaps of synthetic CITE-seq data (supplementary Note I). Native counts represent the ground truth, which are the signals we aimed to recover from observed counts. scAR refers to the version in **Fig. 1**, which we use cell-free droplets to estimate the ambient frequencies. scAR v1 refers to the version in (**a**). **c**, scAR v1 fails to learn the real ambient frequencies. Each dot represents a protein marker. **d**, The noise ratios estimated by two versions of scAR. Each dot represents a cell, colors represent cell type.

**Supplementary Fig. 11.**
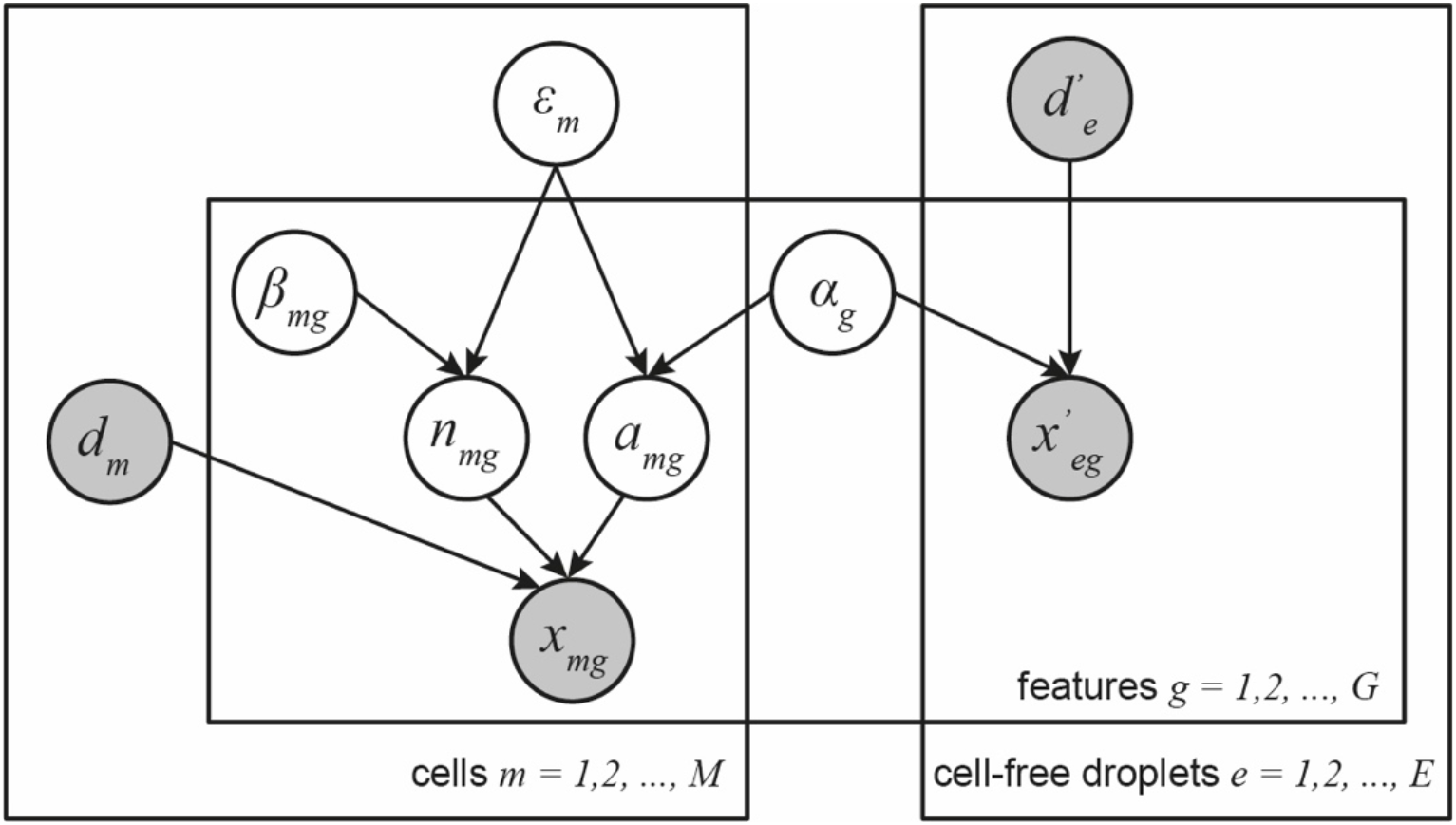
Graphic representation of scAR. The plates (three rectangles) represent independent replications, here, meaning individual cells, features, and cell-free droplets, respectively. Grey circles represent observed random variables, e.g., *d*_*m*_ represents total counts in cell *m* and *x*_*mg*_ represents the observed count of feature *g* in cell *m*. Open circles represent latent random variables. Edges denote conditional dependencies among the variables.

**Supplementary Fig. 12.**
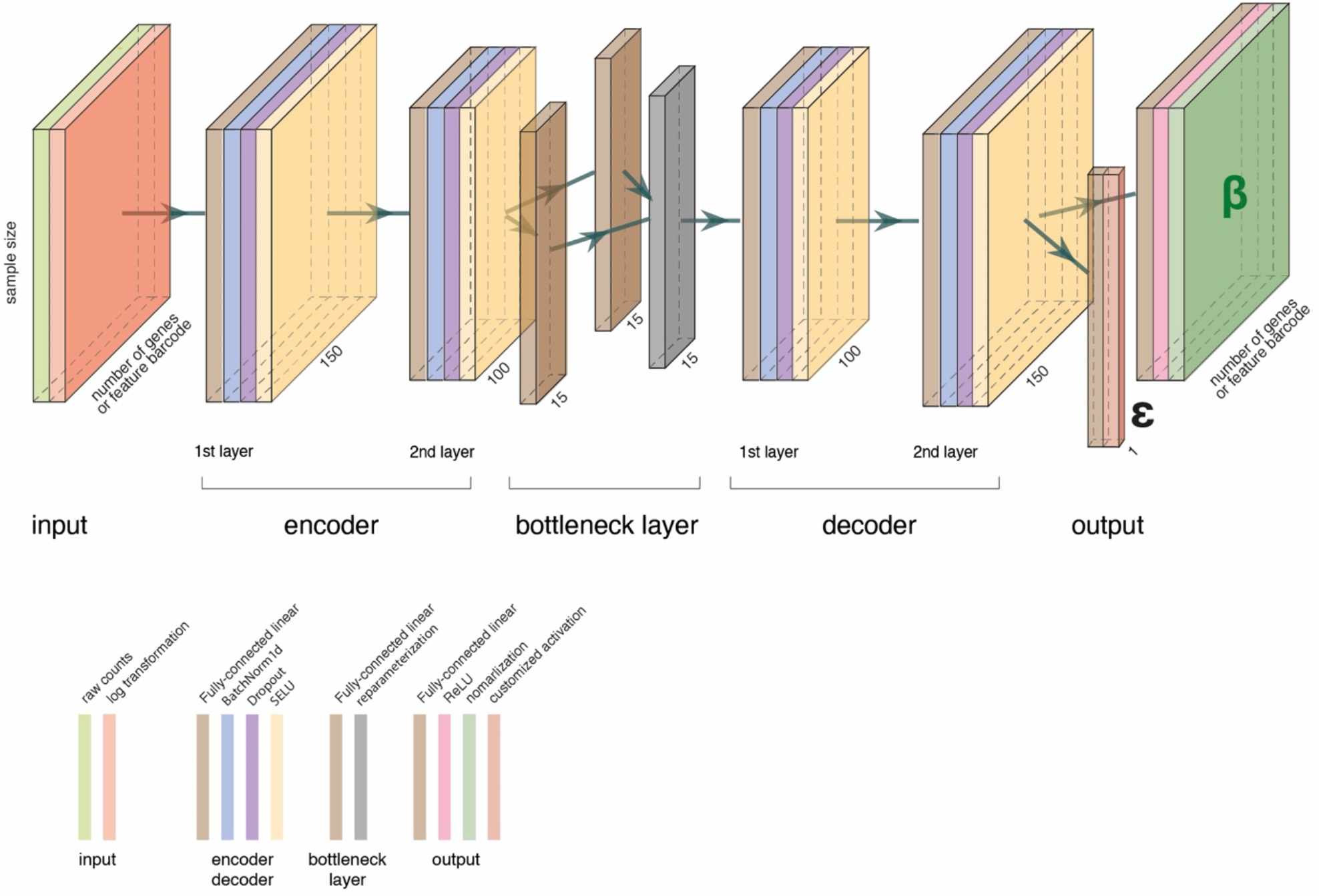
The architecture of VAE in scAR. The optimized dimension numbers of neural network layers are indicated and used as default parameters in scAR. They can also be modified by assigning optional arguments in the scAR command line tool.

**Supplementary Fig. 13.**
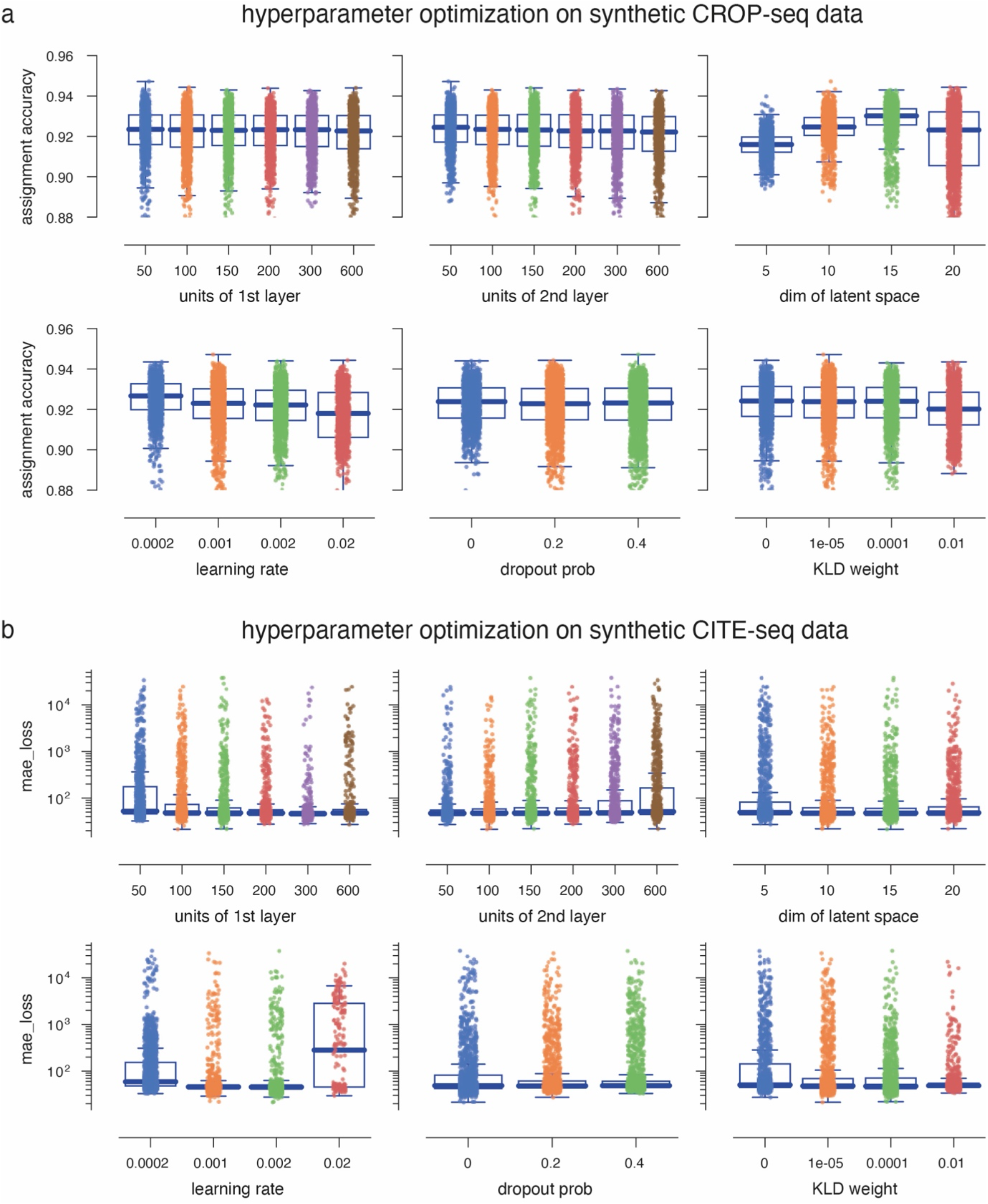
Hyperparameter optimization of scAR. Two synthetic datasets (supplementary Note I), simulating CROP-seq data type and CITE-seq data type, were used to optimize scAR to allow generalized performance. We performed grid search to identify the optimal parameter set. In total, there are 6912 sets of parameters in each dataset. **a**, Hyperparameter optimization on a synthetic CROP-seq dataset. In this class of single-cell omics technologies, assignment of identify barcodes is key information (classification problem), so we used assignment accuracy as a metric to compare performance among parameters. **b**, Hyperparameter optimization on a synthetic CITE-seq dataset. As with scRNAseq, levels of feature barcodes are important information (regression problem). So, we use Mean Absolute Error (MAE loss) as metric in this case.

## Supplementary Note I

### Synthetic CROP-seq and CITE-seq data

For the development and evaluation of scAR, we built a tool to simulate single cell omics data. It is worth noticing that the most recognized package, Splatter^4^, generates synthetic scRNAseq counts with random noise, which assumes that background noise is random. In addition, it is designed to produce scRNAseq data, which is not appropriate to simulate CROP-seq and CITE-seq experiments. To overcome these, we design simulation modules to synthesize CROP-seq and CITE-seq data under ambient signal hypothesis.

#### 1) Simulation of CROP-seq data

The definition of variables in this module is as follows.

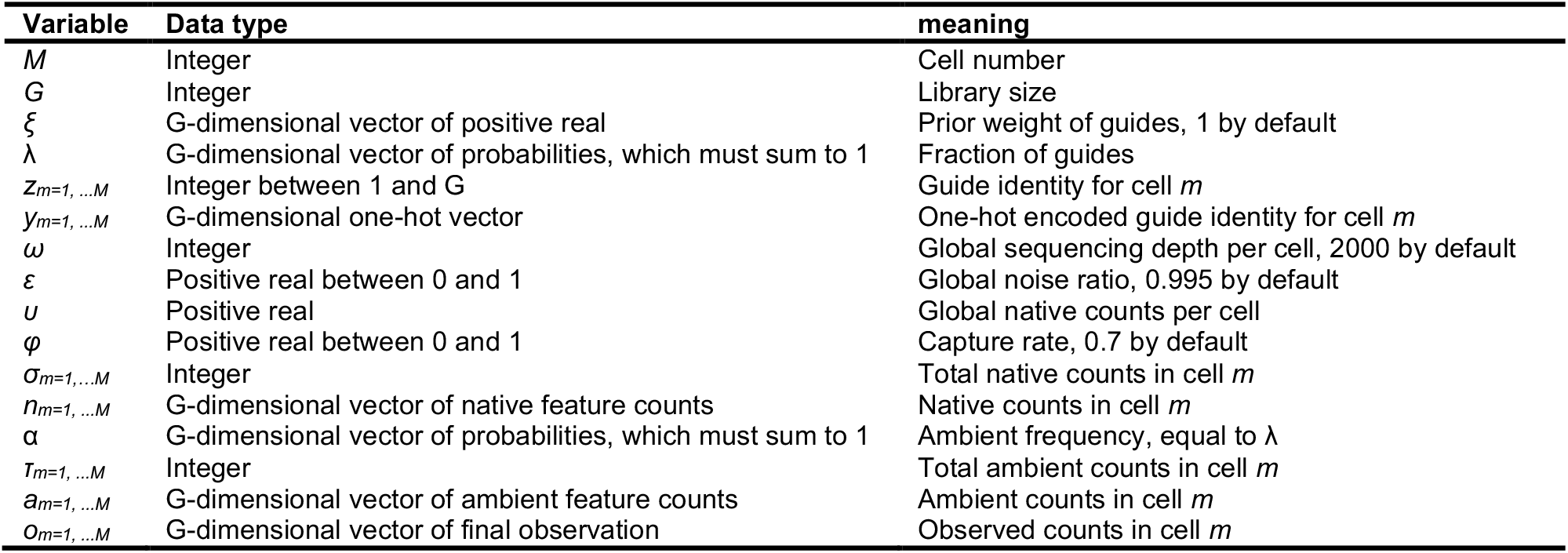

##### Step 1, Generate native signals

We first draw *λ* from a Dirichlet distribution:

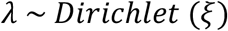

The guide identities then can be sampled from Categorical distribution:

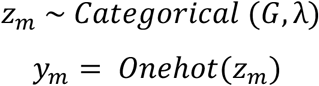

CROP-seq libraries generally use an identical background construct, meaning all guides share an identical promoter. We therefore assume that per cell expression of native signals is drawn from a same distribution, independent of guide sequences. The expected native expression per cell can be calculated as:

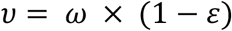

As there are technical dropout events, where there is a certain probability that molecules are not captured by sequencer. To reflect this, we draw the final native counts from a binomial distribution:

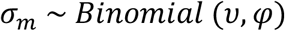

As we assume that each cell integrates only one guide in the case of CROP-seq, we can get the native expression vector for cell *m*:

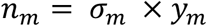

##### Step 2, Generate ambient signals

Under ambient signal hypothesis, the ambient frequency α is correlated with the frequencies of native signals. And the native expression of guide is driven by the same promoter, so we have:

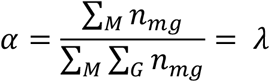

Similarly, when considering the dropout event, we can draw actual ambient counts from a Binomial distribution:

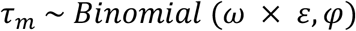

Then, the ambient signals per cell can be drawn from a Multinomial distribution:

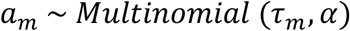

##### Step 3, Sum

All together, we get the final observation,

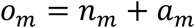

#### 2) Simulation of CITE-seq data

The definition of variables in this module is as follows.

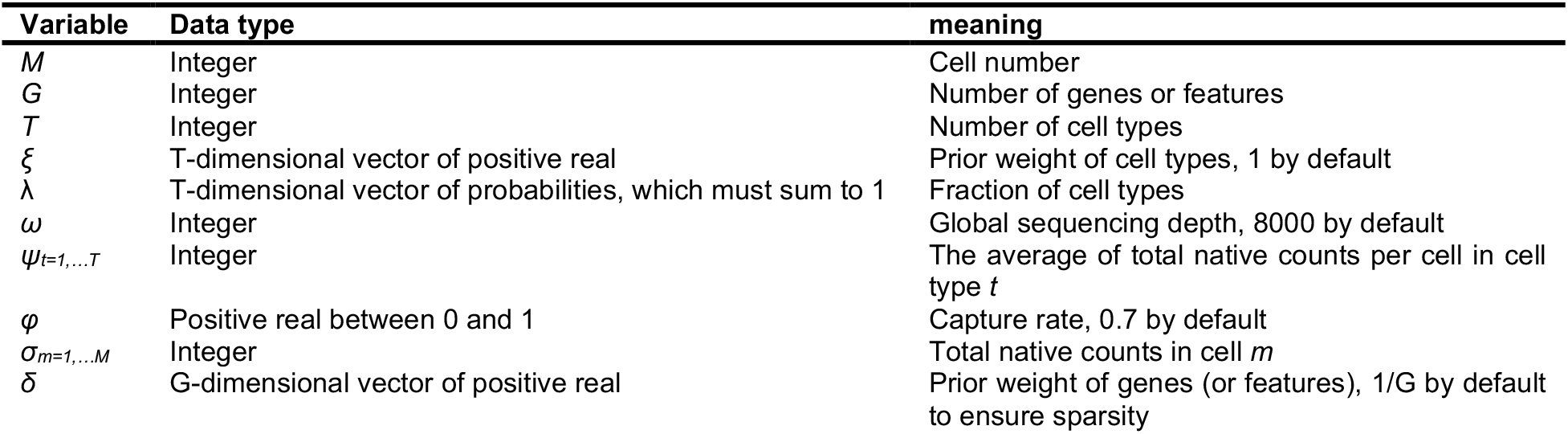
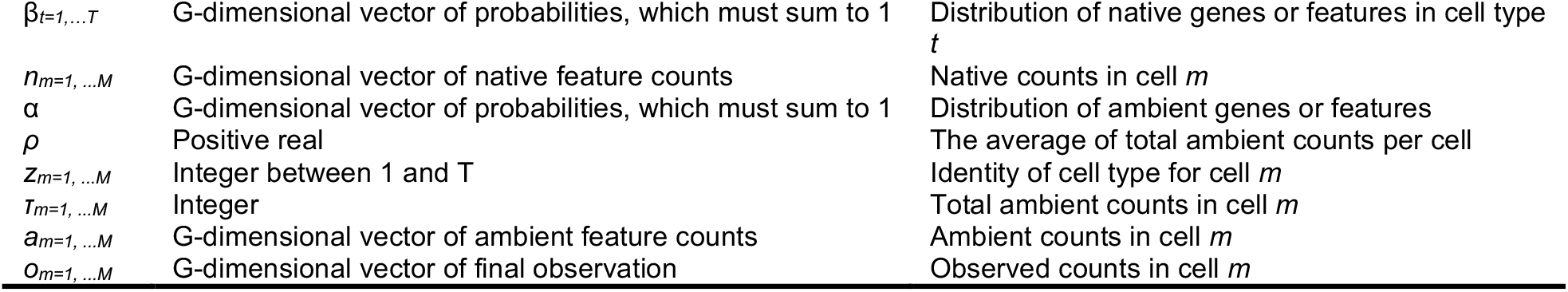

##### Step 1, Generate native signals

We first draw *λ* from a Dirichlet distribution:

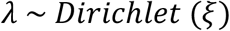

The cell type identities then can be sampled from Categorical distribution:

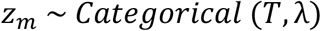

Different cell types generally have different total UMI counts (i.e., sequencing depth) in the case of CITE-seq protein counts, we therefore sample total native UMI counts per cell type from a discrete Uniform distribution:

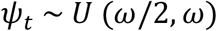

As there are technical dropout events, where there is a certain probability that molecules are not captured by sequencer. To reflect this, we draw the total native counts from a binomial distribution:

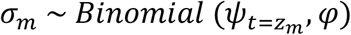

We next sample native expression frequencies (*β*_*t*_) from a Dirichlet distribution:

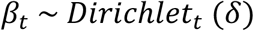

Together, for cell *m*, the native expression at feature level can be drawn from multinomial distribution:

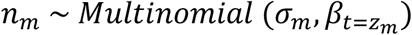

##### Step 2, Generate ambient signals

Under ambient signal hypothesis, the ambient frequency α is correlated with the frequencies of native signals. So, we can calculate α using the following formula:

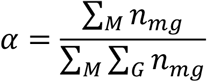

We assume that the total ambient counts are drawn from a discrete Uniform distribution:

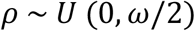

Similarly, when considering the dropout event, we can draw actual ambient counts from a Binomial distribution:

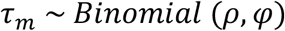

Then, the ambient signals per cell can be drawn from a Multinomial distribution:

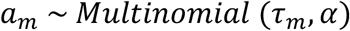

##### Step 3, Sum

All together, we get the final observation,

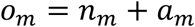

