## Supplementary materials for "Probabilistic machine learning ensures accurate ambient denoising in droplet-based single-cell omics"

### Supplementary Table

| sgRNA number | sgRNA name | Guide sequence | Target genes | Group names | Library name |
|---|---|---|---|---|---|
| 1 | CDH1_g1 | AGTTCCGACGCCACTGAGAG | CDH1 | CDH1 | CROP-seq_pilot |
| 2 | CDH1_g2 | GACTTGCGAGGGACGCATTC | CDH1 | CDH1 | CROP-seq_pilot |
| 3 | CDH1_g3 | CCGAGAGGCTGCGGCTCCAA | CDH1 | CDH1 | CROP-seq_pilot |
| 4 | CDH1_g4 | CGGTGACGACGGGAGAGGAA | CDH1 | CDH1 | CROP-seq_pilot |
| 5 | CDH1_g5 | CCTCAGGACCCGAACTTTCT | CDH1 | CDH1 | CROP-seq_pilot |
| 6 | ESR1_TSS1_g1 | TCCGTTCTGAGTCGGTAGAC | ESR1 | ESR1_TSS1 | CROP-seq_pilot |
| 7 | ESR1_TSS1_g2 | AGCTCTTTAACAGGCTCGAA | ESR1 | ESR1_TSS1 | CROP-seq_pilot |
| 8 | ESR1_TSS1_g3 | CTATAGAATGGGCAGGAGAA | ESR1 | ESR1_TSS1 | CROP-seq_pilot |
| 9 | ESR1_TSS1_g4 | GTATGTTATCTGGAACAGAC | ESR1 | ESR1_TSS1 | CROP-seq_pilot |
| 10 | ESR1_TSS1_g5 | GGTGTGCTGTACTAAGAAAA | ESR1 | ESR1_TSS1 | CROP-seq_pilot |
| 11 | FOXA1_g1 | GCCGCCCGTCGCTTCGCACA | FOXA1 | FOXA1 | CROP-seq_pilot |
| 12 | FOXA1_g2 | CCAACGCCACCCGGGCGAAG | FOXA1 | FOXA1 | CROP-seq_pilot |
| 13 | FOXA1_g3 | CGCCTCCGCGGGAAGTGAGC | FOXA1 | FOXA1 | CROP-seq_pilot |
| 14 | FOXA1_g4 | CAACTGCACTTGCCTCGCAG | FOXA1 | FOXA1 | CROP-seq_pilot |
| 15 | FOXA1_g5 | GTGAGCGGGCTGCCTCTGCG | FOXA1 | FOXA1 | CROP-seq_pilot |
| 16 | GATA3_g1 | CTGTGGCGCGACGCAACTTA | GATA3 | GATA3 | CROP-seq_pilot |
| 17 | GATA3_g2 | GCAACGCAATCTGACCGAGC | GATA3 | GATA3 | CROP-seq_pilot |
| 18 | GATA3_g3 | GCGGCGGCGTACGACCTGCT | GATA3 | GATA3 | CROP-seq_pilot |
| 19 | GATA3_g4 | TTCGCTACCCAGGTTGGTAC | GATA3 | GATA3 | CROP-seq_pilot |
| 20 | GATA3_g5 | TTAGGTCCTCCCAAGTGGTT | GATA3 | GATA3 | CROP-seq_pilot |
| 21 | GRHL2_g1 | ACTAAAGGGTACAAGCCCGA | GRHL2 | GRHL2 | CROP-seq_pilot |
| 22 | GRHL2_g2 | CGCGGAGTCCTCCTGGATCG | GRHL2 | GRHL2 | CROP-seq_pilot |
| 23 | GRHL2_g3 | CCTCACCTAGCCGGAAAGGT | GRHL2 | GRHL2 | CROP-seq_pilot |
| 24 | GRHL2_g4 | GTGTGTGAGAGCGCCCGAGA | GRHL2 | GRHL2 | CROP-seq_pilot |
| 25 | GRHL2_g5 | CCTTGCGAGAAAGTTACCTG | GRHL2 | GRHL2 | CROP-seq_pilot |
| 26 | KMT2D_g1 | AACAGACGAGATGCCTCCGG | KMT2D | KMT2D | CROP-seq_pilot |
| 27 | KMT2D_g2 | GATAGAGGCGTCTCAAGTGC | KMT2D | KMT2D | CROP-seq_pilot |
| 28 | KMT2D_g3 | GACAAGGGCGACTCCTCCAG | KMT2D | KMT2D | CROP-seq_pilot |
| 29 | KMT2D_g4 | GGGCAATTCCTCAGGTGGCG | KMT2D | KMT2D | CROP-seq_pilot |
| 30 | KMT2D_g5 | GGGCGATGCTTCAGGTGGTG | KMT2D | KMT2D | CROP-seq_pilot |
| 31 | KMT2C_g1 | GACTAGGATGTCGTCGGAGG | KMT2C | KMT2C | CROP-seq_pilot |
| 32 | KMT2C_g2 | CGCACTCACACACATCGGCG | KMT2C | KMT2C | CROP-seq_pilot |
| 33 | KMT2C_g3 | GGATCCCGGTCCTCCTCCTG | KMT2C | KMT2C | CROP-seq_pilot |
| 34 | KMT2C_g4 | AAATGCGAGAGGCTGAGCCG | KMT2C | KMT2C | CROP-seq_pilot |
| 35 | KMT2C_g5 | TCTCGCATTTCCCGCAGCCC | KMT2C | KMT2C | CROP-seq_pilot |
| 36 | CTRL_g1 | TCTCGTCTGATACCTCGGTC | OR2L13 | CTL | CROP-seq_pilot |
| 37 | CTRL_g2 | CTCATCGTGGTCGGCGGTCG | OR2L13 | CTL | CROP-seq_pilot |
| 38 | CTRL_g3 | GCGGCGTCTTTGGCAGTAGT | OR2L13 | CTL | CROP-seq_pilot |
| 39 | CTRL_g4 | GGCGTGCTTGCGGGTCCAGG | OR2L13 | CTL | CROP-seq_pilot |
| 40 | CTRL_g5 | CGCTGCTGCGAGACCAGCCG | OR2L13 | CTL | CROP-seq_pilot |
| 41 | CTRL_g6 | ACTCACCTCAACCGTATGGA | CTL | CTL | CROP-seq_pilot |
| 42 | CTRL_g7 | CTGCAAGTAACCCATGCACC | CTL | CTL | CROP-seq_pilot |
| 43 | CTRL_g8 | ATGCACTCAGCAAGTCTAAC | CTL | CTL | CROP-seq_pilot |
| 44 | CTRL_g9 | GGCTGTGAAGAACCAGAAGT | CTL | CTL | CROP-seq_pilot |
| 45 | CTRL_g10 | GCTGCCTGTCCTTTGAGTCA | CTL | CTL | CROP-seq_pilot |
| 46 | CTRL_g11 | CCGCAGCAATATCTTGGCTC | CTL | CTL | CROP-seq_pilot |
| 47 | CTRL_g12 | GGGCTCTCCAACTCACCAGG | CTL | CTL | CROP-seq_pilot |
| 48 | CTRL_g13 | TGCTCAGCAGACTAGGCAGC | CTL | CTL | CROP-seq_pilot |
| 49 | CTRL_g14 | GAAGCTCTGCTCAGCAGACT | CTL | CTL | CROP-seq_pilot |
| 50 | CTRL_g15 | TCTGTCTCTGAGCTAGACTT | CTL | CTL | CROP-seq_pilot |
| 51 | TRPS1_g1 | GACGTAATGCGCGGAGACTG | TRPS1 | TRPS1 | CROP-seq_pilot |
| 52 | TRPS1_g2 | CTTGAAACTGACGTAATGCG | TRPS1 | TRPS1 | CROP-seq_pilot |
| 53 | TRPS1_g3 | AGAGCAATCGAGAGGACGCG | TRPS1 | TRPS1 | CROP-seq_pilot |
| 54 | TRPS1_g4 | AAGGCGAGAGAGCAATCGAG | TRPS1 | TRPS1 | CROP-seq_pilot |
| 55 | TRPS1_g5 | GGATGTGCCCGGTGCCGGGT | TRPS1 | TRPS1 | CROP-seq_pilot |
| 56 | YAP1_g1 | CCGCCAGACCAGTGGAGCCG | YAP1 | YAP1 | CROP-seq_pilot |
| 57 | YAP1_g2 | CCTCCGTCAAGGGAGTTGGA | YAP1 | YAP1 | CROP-seq_pilot |
| 58 | YAP1_g3 | CGGCGCTGTCCTCGCTCTCA | YAP1 | YAP1 | CROP-seq_pilot |
| 59 | YAP1_g4 | GGCGAGTTTCTGTCTCAGTC | YAP1 | YAP1 | CROP-seq_pilot |
| 60 | YAP1_g5 | CTGCGAGGCACTCGGACCTG | YAP1 | YAP1 | CROP-seq_pilot |
| 61 | CTL _g16 | TAGATCTGAAAGGCTGGGAT | CTL | CTL | CROP-seq_pilot |
| 62 | CTL _g17 | TGTCTCCTACTGCGTGTTGA | CTL | CTL | CROP-seq_pilot |
| 63 | CTL _g18 | TCTTAATGATAGAATCTTCC | CTL | CTL | CROP-seq_pilot |
| 64 | CTL _g19 | GCTCCCAGTGTCCTGTGATA | CTL | CTL | CROP-seq_pilot |
| 65 | CTL _g20 | AAGCACCCAGTAGTAAAACA | CTL | CTL | CROP-seq_pilot |

| | | | | | |
|---|---|---|---|---|---|
| 66 | CTRL_g21 | CTGAAAAAGGAAGGAGTTGA | CTL | CTL | CROP-seq_pilot |
| 67 | CTRL_g22 | AAGATGAAAGGAAAGGCGTT | CTL | CTL | CROP-seq_pilot |
| 68 | CTRL_g23 | TGCGCGGCTTGGGAAGCCCA | CTL | CTL | CROP-seq_pilot |
| 69 | CTRL_g24 | GACGCGAGGAAGGAGGGCGC | CTL | CTL | CROP-seq_pilot |
| 70 | enh588_g1 | GGATCTGCAGGCCCAAGGTC | CCND1 | enh588 | CROP-seq_pilot |
| 71 | enh588_g2 | CTCTCAGTCATCCTTGACCTT | CCND1 | enh588 | CROP-seq_pilot |
| 72 | enh588_g3 | GCTCTCAGTCATCCCTGACCT | CCND1 | enh588 | CROP-seq_pilot |
| 73 | enh588_g4 | TCCTCTAGCAGACGGCCCTG | CCND1 | enh588 | CROP-seq_pilot |
| 74 | enh588_g5 | TCTGCTAGAGGATCACTCCT | CCND1 | enh588 | CROP-seq_pilot |
| 75 | enh588_g6 | GGCGGAGTCATGCCAGCTCA | CCND1 | enh588 | CROP-seq_pilot |
| 76 | CCND1_g1 | GCAGCAGAGTCCGCACGCTC | CCND1 | CCND1 | CROP-seq_pilot |
| 77 | CCND1_g2 | GGTGAGTAGCAAAGAAACGT | CCND1 | CCND1 | CROP-seq_pilot |
| 78 | CCND1_g3 | ACTCCGCCGCAGGGCAGGCG | CCND1 | CCND1 | CROP-seq_pilot |
| 79 | CCND1_g4 | CTATGAAAACCGGACTACAG | CCND1 | CCND1 | CROP-seq_pilot |
| 80 | ESR1_TSS2_g1 | AAGCCGGGCGACCCGAC | ESR1 | ESR1_TSS2 | CROP-seq_pilot |
| 81 | ESR1_TSS2_g2 | GGCGCACGAGGATCTGCTAA | ESR1 | ESR1_TSS2 | CROP-seq_pilot |
| 82 | ESR1_TSS2_g3 | GGAGCCCAGGAGCTGGCGGA | ESR1 | ESR1_TSS2 | CROP-seq_pilot |
| 83 | ESR1_TSS2_g4 | TCAGGGCAAGGCAACAGTCCC | ESR1 | ESR1_TSS2 | CROP-seq_pilot |
| 84 | ESR1_TSS2_g5 | GGAGACCAGTACTTAAAGT | ESR1 | ESR1_TSS2 | CROP-seq_pilot |
| 85 | TP53_g1 | ATGAGTCCTCTCTGAGTCAC | TP53 | TP53 | CROP-seq_pilot |
| 86 | TP53_g2 | TCAGGAGCTTACCCAATCCA | TP53 | TP53 | CROP-seq_pilot |
| 87 | TP53_g3 | CCGAGAGCCCGTGACTCAGAG | TP53 | TP53 | CROP-seq_pilot |
| 88 | TP53_g4 | TGGGGACTTAGCGAGTTT | TP53 | TP53 | CROP-seq_pilot |
| 89 | TP53_g5 | GGAAGCGTGTCACCGTCG | TP53 | TP53 | CROP-seq_pilot |
| 90 | PIK3CA_g1 | TCTCCCAGCGTCGGCCCG | PIK3CA | PIK3CA | CROP-seq_pilot |
| 91 | PIK3CA_g2 | AGCGTGAGTAGAGCGCGGAC | PIK3CA | PIK3CA | CROP-seq_pilot |
| 92 | PIK3CA_g3 | GAGAGGGTGCGGCGATCGC | PIK3CA | PIK3CA | CROP-seq_pilot |
| 93 | PIK3CA_g4 | CCCCGAGCGTGAGTAGAGCG | PIK3CA | PIK3CA | CROP-seq_pilot |
| 94 | PIK3CA_g5 | GGAGTCTCCGGCACCCACC | PIK3CA | PIK3CA | CROP-seq_pilot |
| 95 | SUZ12_g1 | GGGCCGCCCGGCGGGTAGCTGG | SUZ12 | SUZ12 | CROP-seq_pilot |
| 96 | SUZ12_g2 | CTCCGGCGGACCGAGGGGGGA | SUZ12 | SUZ12 | CROP-seq_pilot |
| 97 | SUZ12_g3 | CGGAGCGAGGCCAGGGTA | SUZ12 | SUZ12 | CROP-seq_pilot |
| 98 | SUZ12_g4 | CAGGCTCCGGCGGACCGAGG | SUZ12 | SUZ12 | CROP-seq_pilot |
| 99 | SUZ12_g5 | TATTGCAGGCGCTTGCTCTC | SUZ12 | SUZ12 | CROP-seq_pilot |

**Supplementary Table 1 | CROP-seq libraries.**

#### Supplementary Figures

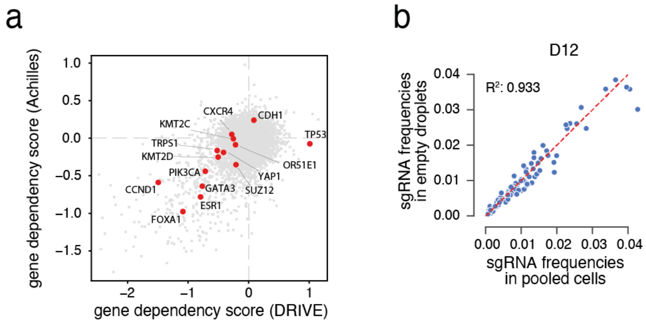


**Supplementary Fig. 1 | Validation of CROP-seq libraries. a**, The gene dependency scores of most target genes show negative effects on the cell growth of MCF7. One exception is the tumour repressor TP53, down-regulation of which promotes tumour growth. Data are originally from two pooled screening projects (DRIVE and Achilles) and obtained from https://depmap.org/. **b,** Scatterplot of sgRNA frequencies between two independent experiments. The X-axis represents sgRNA frequencies in bulk DNA sequencing, Y-axis represents sgRNA frequencies obtained by averaging RNA counts in cell-free droplets from 10x scRNAseq. Each dot represents an sgRNA, the red dashed lines represent y=x, and coefficients of determination ($R^2$ scores) is shown. D12 sample was shown

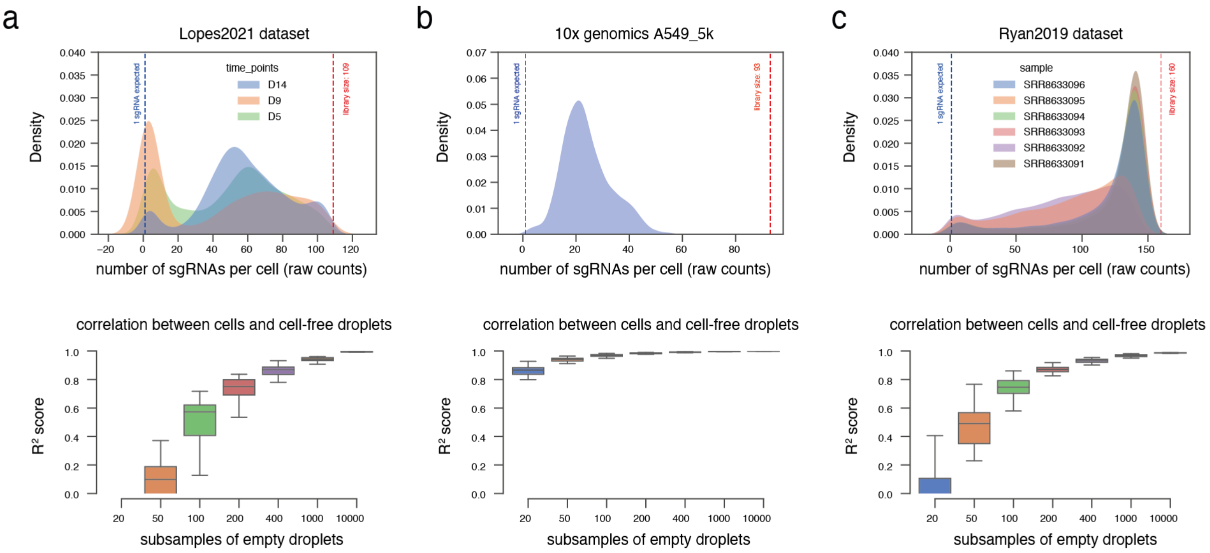


**Supplementary Fig. 2 | supplementary to Fig. 2, ambient noise widely exists in single-cell omics datasets.** Ambient contamination is observed in several public datasets and the background noise is highly correlated with endogenous signals in cells. **a,** The Lopes2021 dataset[1]. **b,** The A549_5k dataset from 10x genomics[2]. **c,** The Ryan2019 dataset[3].

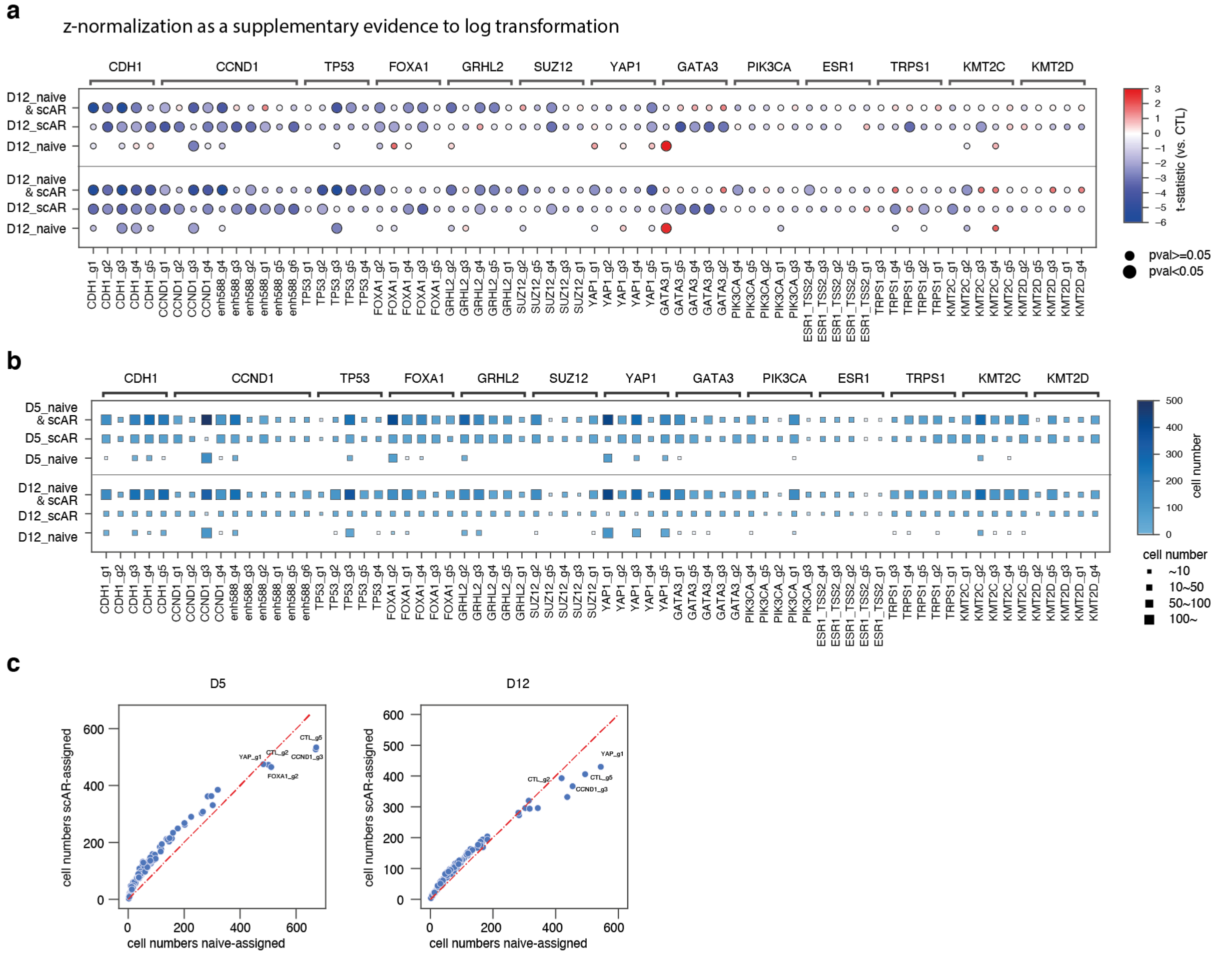


**Supplementary Fig. 3 | Supplementary to Fig. 3.** Evaluation of scAR on CROP-seq. **a,** Similar to **Fig. 3g**, the dotplot shows the overall comparison between two assignment approaches on t-statics of z-normalized expression. The X-axis represents guide groups. Y-axis represents subgroups of cells as exemplified in **Fig. 3e** and **Fig. 3f**, separated by two time points and assignment approaches. Target genes are shown on the top. Their expression (log-transformed) in each group is compared with that in the CTL group and resulting t-statistics are shown by the dot colour. Blue colour indicates down-regulation, and red indicates up-regulation. CTL group is centred at zero. The bimodal sizes of circles represent the p-values from the t-test (the bigger means p<0.05, the smaller means p>=0.05). The asterisk (*) highlights the guide groups where scAR significantly improves the accuracy and hashes (#) mark the groups where scAR underperforms naïve assignment. **b,** The comparison between two assignment approaches on cell number after assignment. Sizes of squares represent the cell numbers of each assignment. **c,** The overall comparisons of cell numbers. Each dot represents an sgRNA.

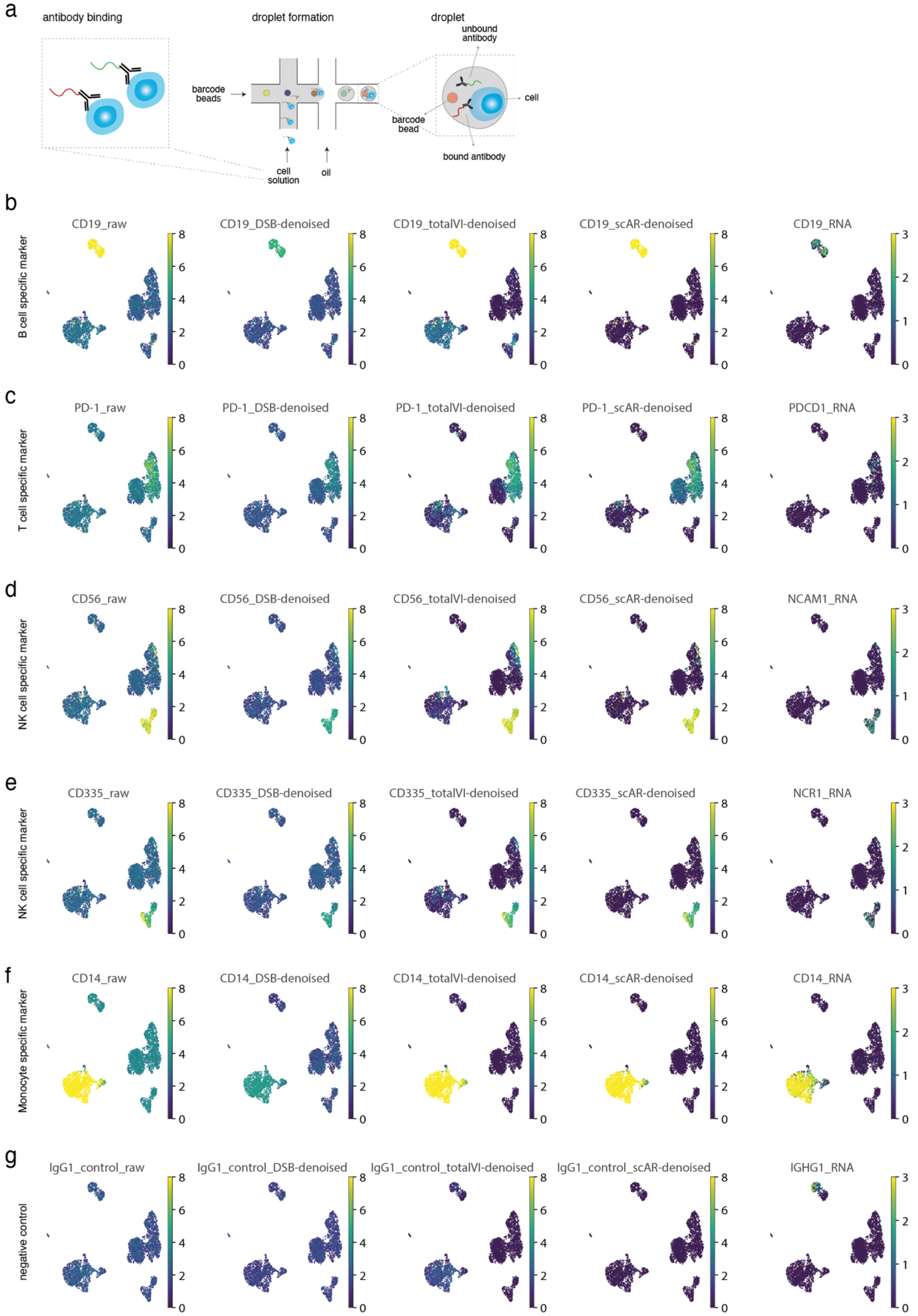

a
antibody binding
droplet formation
droplet
unbound antibody
barcode beads
cell
barcode bead
bound antibody
cell solution
oil
b
B cell specific marker
CD19_raw
CD19_DSB-denoised
CD19_totalVI-denoised
CD19_scAR-denoised
CD19_RNA
c
T cell specific marker
PD-1_raw
PD-1_DSB-denoised
PD-1_totalVI-denoised
PD-1_scAR-denoised
PDCD1_RNA
d
NK cell specific marker
CD56_raw
CD56_DSB-denoised
CD56_totalVI-denoised
CD56_scAR-denoised
NCAM1_RNA
e
NK cell specific marker
CD335_raw
CD335_DSB-denoised
CD335_totalVI-denoised
CD335_scAR-denoised
NCR1_RNA
f
Monocyte specific marker
CD14_raw
CD14_DSB-denoised
CD14_totalVI-denoised
CD14_scAR-denoised
CD14_RNA
g
negative control
IgG1_control_raw
IgG1_control_DSB-denoised
IgG1_control_totalVI-denoised
IgG1_control_scAR-denoised
IGHG1_RNA

**Supplementary Fig. 4 | scAR reduces the non-specific ambient antibodies. a,** The illustration of CITE-seq experiment and the potential source of background noise. **b-g,** UMAPs visualize raw protein counts, DSB-normalised, totalVI-denoised, and scAR-denoised protein counts, and the corresponding RNA counts of cell-specific markers, including a B cell-specific marker CD19 (**b**), a T cell-specific marker PD-1 (**c**), two NK cell-specific markers CD56 (**d**) and CD335 (**e**), and a monocyte-specific marker CD14(**f**), as well as a negative control IgG1 (**g**).

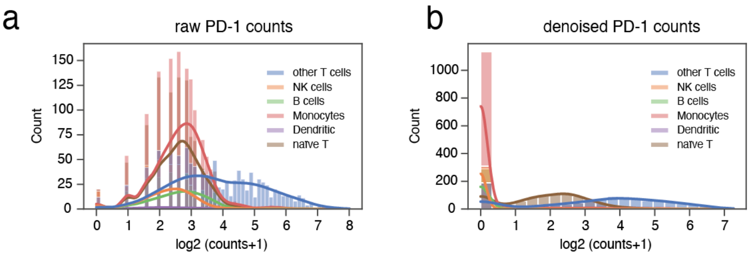


**Supplementary Fig. 5 | scAR removed the non-specific ambient PD-1 but preserved specific PD-1 even when they had similar counts. a,** The distributions of raw PD-1 counts were similar/highly overlapped between cell types. **b,** scAR preserved true PD-1 signal in naïve T cells but removed false PD-1 signal in other cell types.

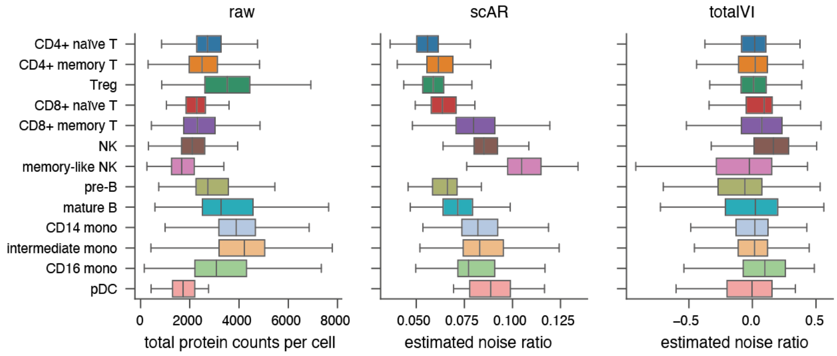


**Supplementary Fig. 6 | scAR removed <10% ambient counts across cell types and showed high consistency across cells. a,** total raw counts of antibodies in cell subtypes. **b,** scAR-estimated noise ratio in cell subtypes. **c,** totalVI-estimated noise ratio in cell subtypes.

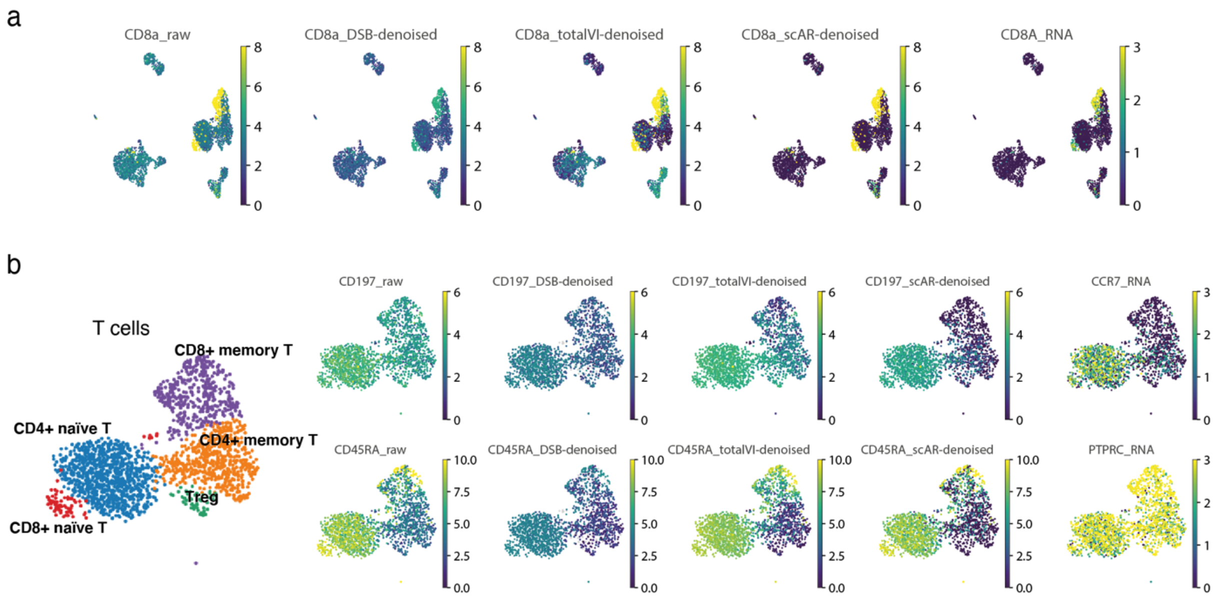


**Supplementary Fig. 7 | scAR denoised proteins show better separation of cell subtypes. a,** scAR-denoised CD8a was exclusively expressed in CD8+ T cells. **b,** scAR-denoised CD197 and CD45RA separated better memory and naïve T cells.

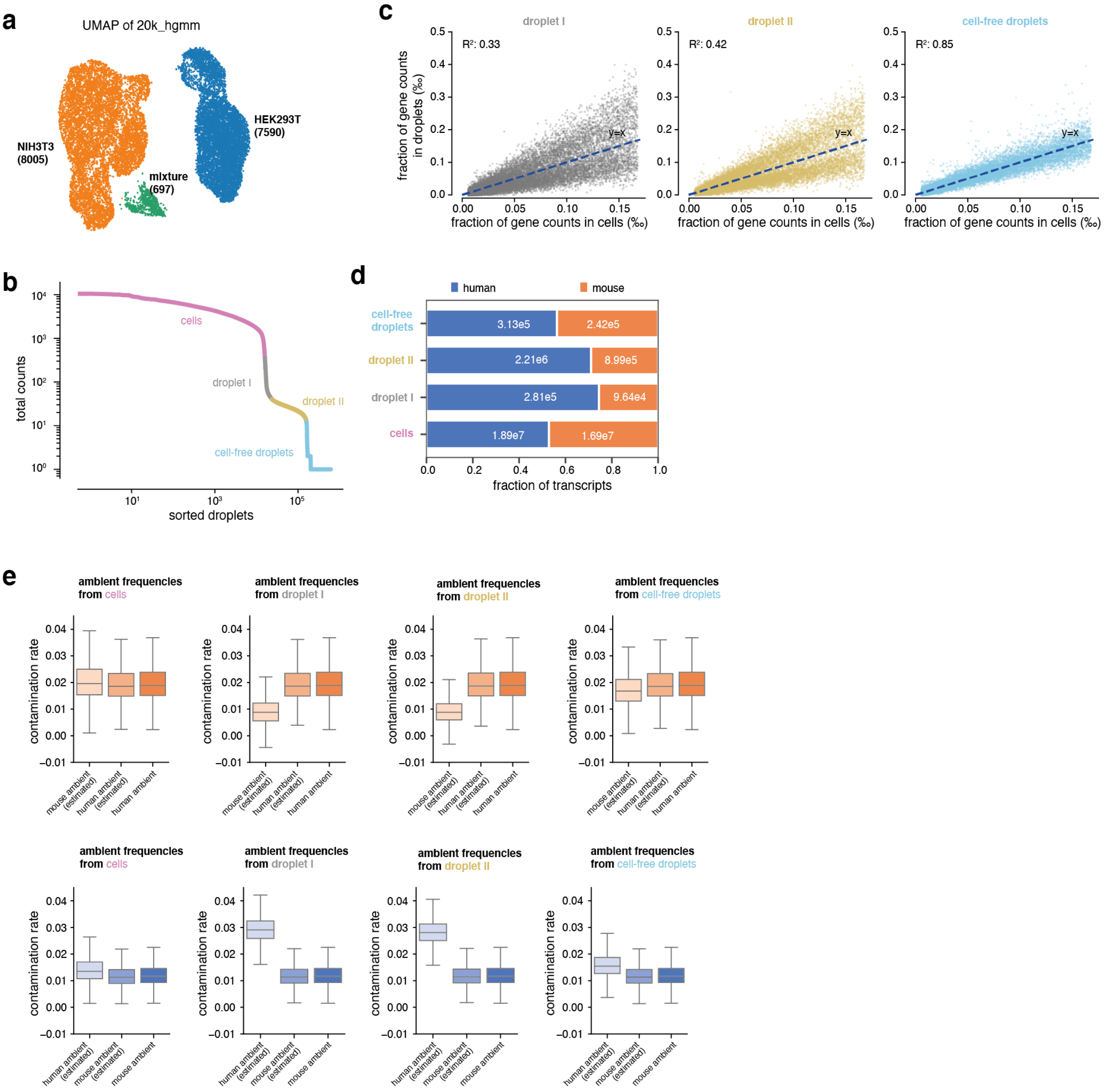


**Supplementary Fig. 8 | Supplementary to Fig. 5.** A public scRNAseq dataset of mixed human HEK293T and mouse NIH3T3 cells (1:1) was selected to demonstrate scAR's ability in noise reduction in transcriptome data. **a,** UMAP shows three populations of cell-containing droplets, HEK293T, NIH3T3 and multiplets. **b,** The kneeplot shows subpopulations of droplets. **c,** Correlation of gene frequencies between subpopulations of droplets and cell-containing droplets. **d,** Fraction of transcripts in subpopulations of droplets. **e,** Boxplots show the percentage of ambient signal in NIH3T3 cells (the upper panel) and HEK293T cells (the bottom panel). Columns separate the different inputted ambient frequencies which were calculated using different subpopulations of droplets. The Y-axis represents contamination ratio, the X-axis represent scAR-estimated inter-species, scAR-estimated cross-species, and true cross-species contamination, respectively. Given that the global ratio of human and mouse transcripts is ~1.11 in cells (**d**), it is reasonable to expect a similar inter-species and cross-species contamination ratios. However, droplet I and II lead to too high estimation of human-sourced contamination, as much as ~3x of mouse source. This may be explained by the over-representation of human transcripts in droplet I and II (**d**). The higher human transcripts in ambient frequencies as input, the more counts to be identified as background noise by scAR. The best estimate of ambient frequencies should be drawn from the population of cell-free droplets, as the estimated noise ratios are in a reasonable range in both cell lines – ambient signals from human sources are slightly stronger than mouse sources in both cell lines. These observations also suggest that compositions in these droplets are clearly different, e.g., droplet I and II may contain more human cell debris.

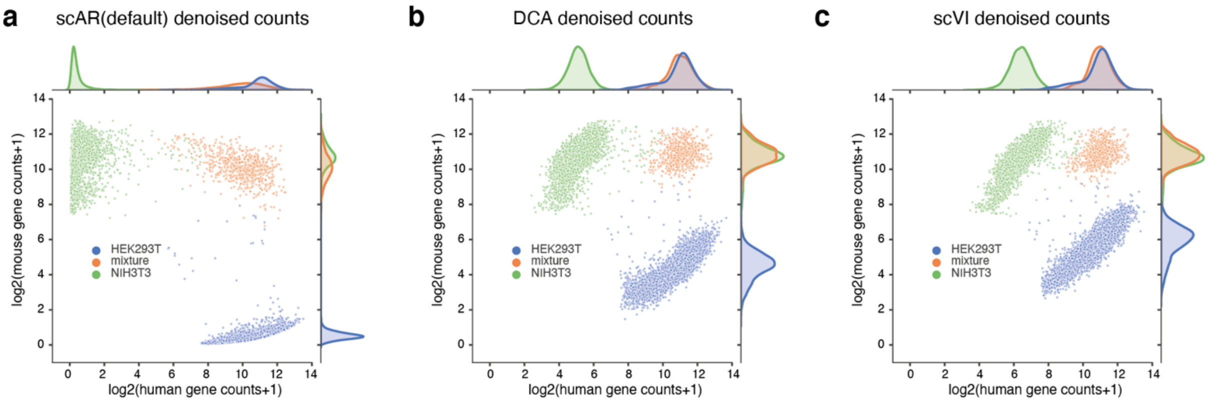


**Supplementary Fig. 9 | Supplementary to Fig. 5. Benchmarking of mRNA count denoising methods. a,** scAR denoised mRNA counts under the default setting – ambient frequencies were calculated using cells. **b,** DCA denoised mRNA counts under its default setting (epochs and batch size were aligned to scAR at epochs = 800, batch_size = 64). **c,** scVI denoised mRNA counts under its default setting (epochs and batch size were aligned to scAR at epochs = 800, batch_size = 64).

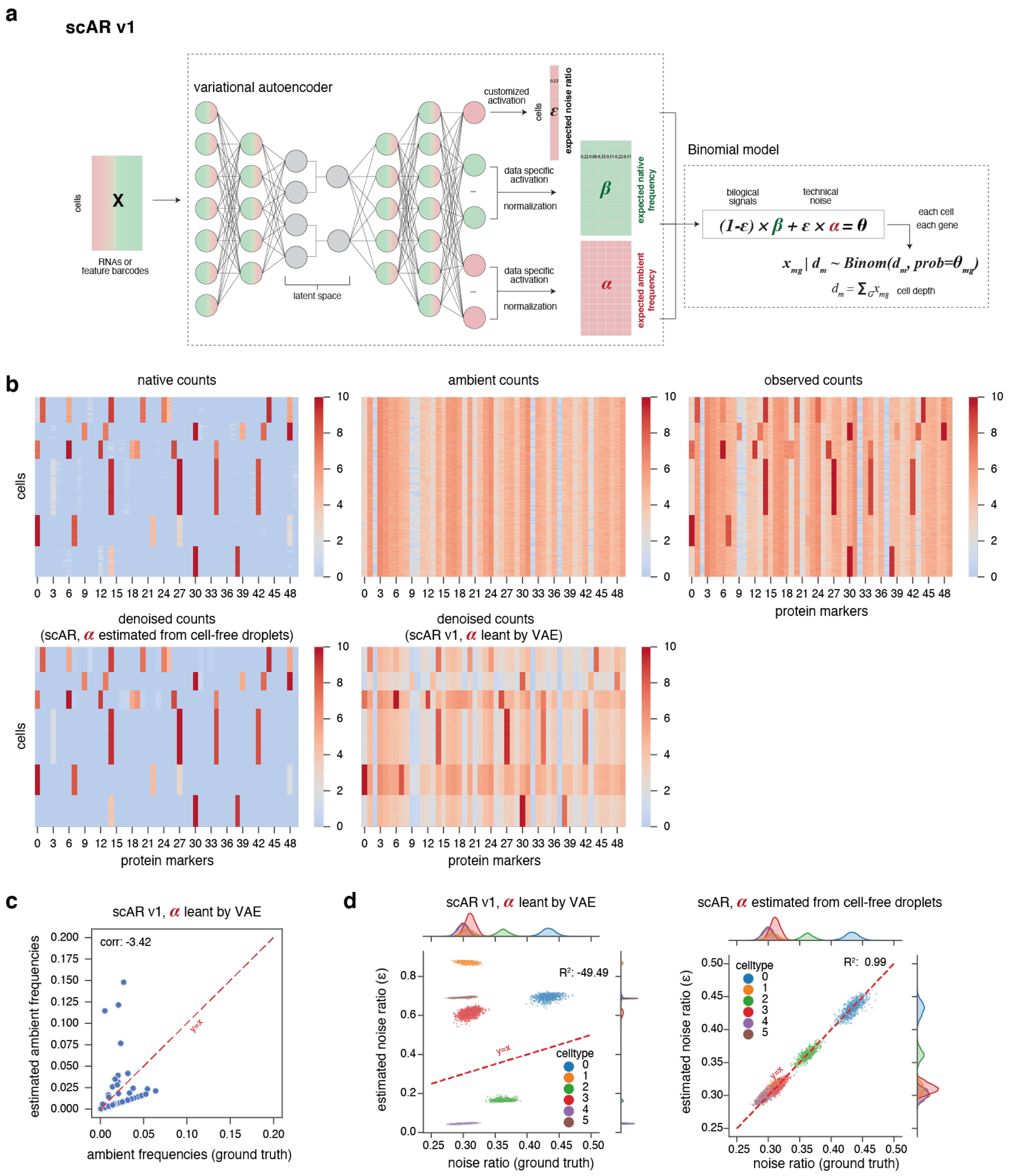


**Supplementary Fig. 10 | The performance of two versions of scAR.** We compared two versions of scAR to demonstrate the necessary of using cell-free droplets. **a,** scAR v1 is an early version of scAR in which we fully relied on VAE to learn the ambient frequencies. **b,** Heatmaps of synthetic CITE-seq data (supplementary Note I). Native counts represent the ground truth, which are the signals we aimed to recover from observed counts. scAR refers to the version in **Fig. 1**, which we use cell-free droplets to estimate the ambient frequencies. scAR v1 refers to the version in (**a**). **c,** scAR v1 fails to learn the real ambient frequencies. Each dot represents a protein marker. **d,** The noise ratios estimated by two versions of scAR. Each dot represents a cell, colors represent cell type.

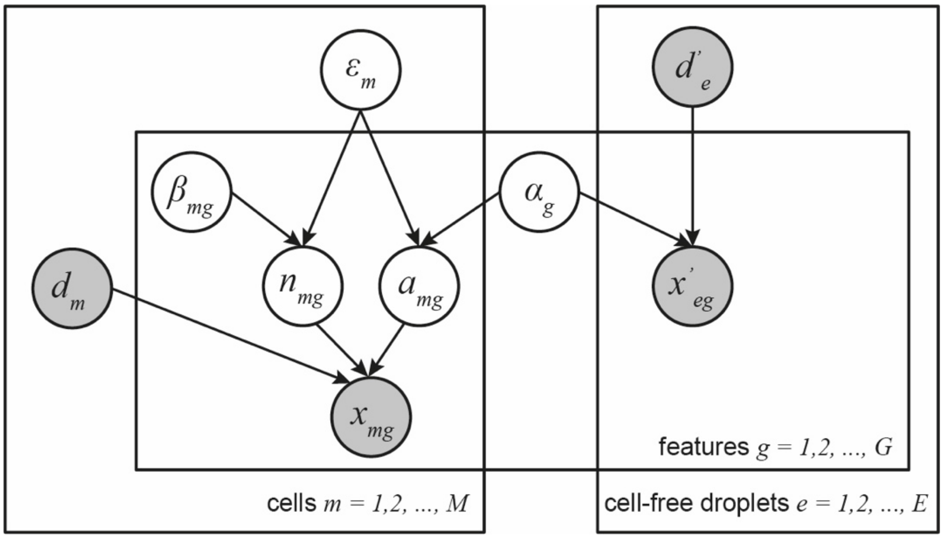


**Supplementary Fig. 11 | Graphic representation of scAR.** The plates (three rectangles) represent independent replications, here, meaning individual cells, features, and cell-free droplets, respectively. Grey circles represent observed random variables, e.g., $d_m$ represents total counts in cell $m$ and $x_{mg}$ represents the observed count of feature $g$ in cell $m$. Open circles represent latent random variables. Edges denote conditional dependencies among the variables.

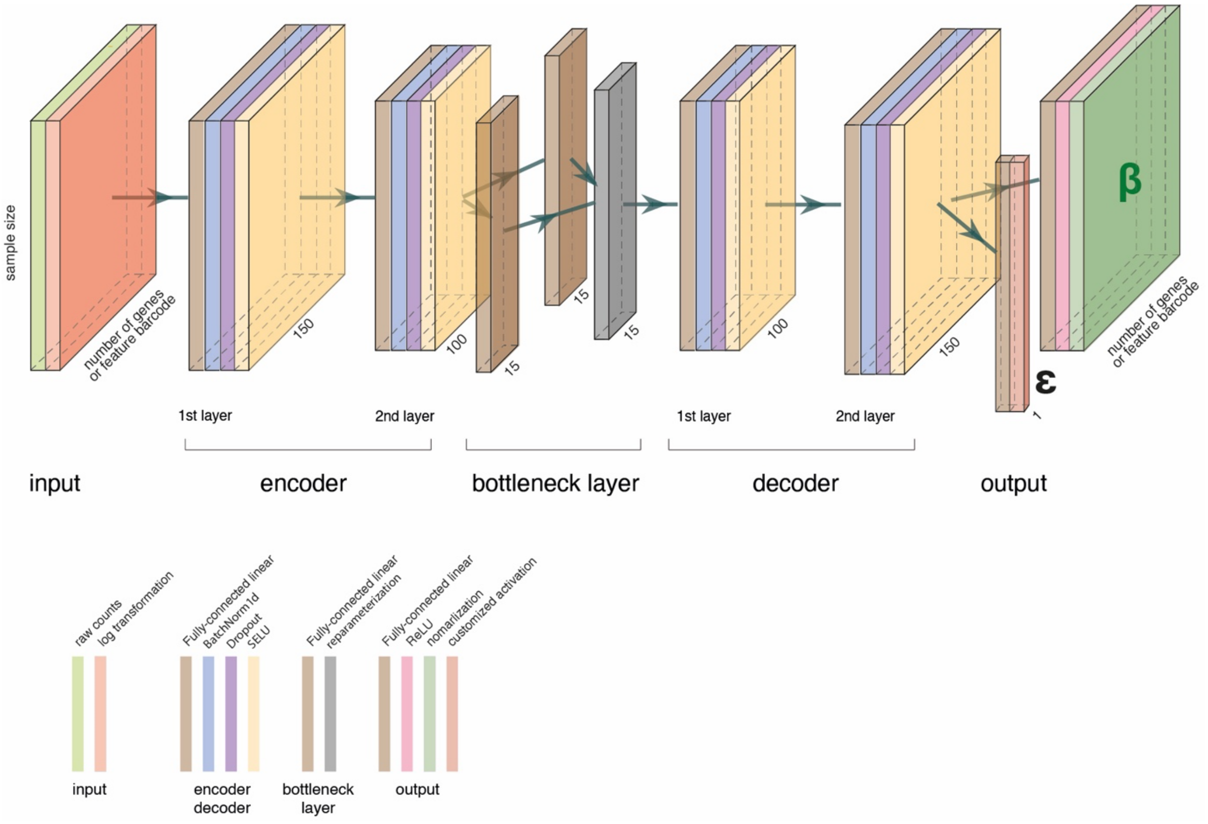


**Supplementary Fig.12 | The architecture of VAE in scAR.** The optimized dimension numbers of neural network layers are indicated and used as default parameters in scAR. They can also be modified by assigning optional arguments in the scAR command line tool.

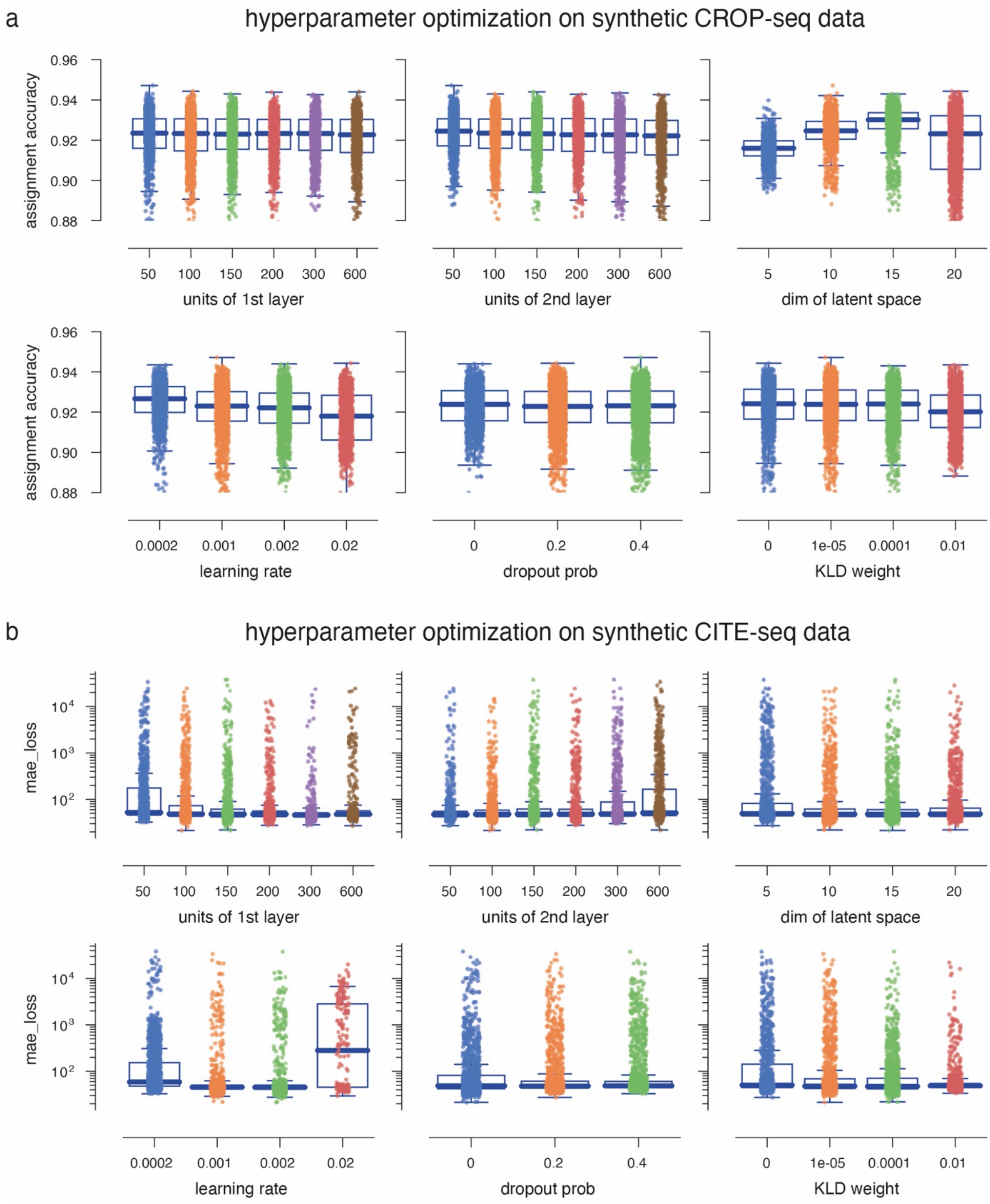


**Supplementary Fig. 13 | Hyperparameter optimization of scAR.** Two synthetic datasets (supplementary Note I), simulating CROP-seq data type and CITE-seq data type, were used to optimize scAR to allow generalized performance. We performed grid search to identify the optimal parameter set. In total, there are 6912 sets of parameters in each dataset. **a,** Hyperparameter optimization on a synthetic CROP-seq dataset. In this class of single-cell omics technologies, assignment of identify barcodes is key information (classification problem), so we used assignment accuracy as a metric to compare performance among parameters. **b,** Hyperparameter optimization on a synthetic CITE-seq dataset. As with scRNAseq, levels of feature barcodes are important information (regression problem). So, we use Mean Absolute Error (MAE loss) as metric in this case.

#### Supplementary Note I

##### *Synthetic CROP-seq and CITE-seq data*

For the development and evaluation of scAR, we built a tool to simulate single cell omics data. It is worth noticing that the most recognized package, Splatter[4], generates synthetic scRNAseq counts with random noise, which assumes that background noise is random. In addition, it is designed to produce scRNAseq data, which is not appropriate to simulate CROP-seq and CITE-seq experiments. To overcome these, we design simulation modules to synthesize CROP-seq and CITE-seq data under ambient signal hypothesis.

1) Simulation of CROP-seq data

The definition of variables in this module is as follows.

| Variable | Data type | meaning |
|---|---|---|
| $M$ | Integer | Cell number |
| $G$ | Integer | Library size |
| $\xi$ | G-dimensional vector of positive real | Prior weight of guides, 1 by default |
| $\lambda$ | G-dimensional vector of probabilities, which must sum to 1 | Fraction of guides |
| $z_{m=1,\ldots M}$ | Integer between 1 and G | Guide identity for cell *m* |
| $y_{m=1,\ldots M}$ | G-dimensional one-hot vector | One-hot encoded guide identity for cell *m* |
| $\omega$ | Integer | Global sequencing depth per cell, 2000 by default |
| $\varepsilon$ | Positive real between 0 and 1 | Global noise ratio, 0.995 by default |
| $\upsilon$ | Positive real | Global native counts per cell |
| $\varphi$ | Positive real between 0 and 1 | Capture rate, 0.7 by default |
| $\sigma_{m=1,\ldots M}$ | Integer | Total native counts in cell *m* |
| $n_{m=1,\ldots M}$ | G-dimensional vector of native feature counts | Native counts in cell *m* |
| $\alpha$ | G-dimensional vector of probabilities, which must sum to 1 | Ambient frequency, equal to λ |
| $\tau_{m=1,\ldots M}$ | Integer | Total ambient counts in cell *m* |
| $a_{m=1,\ldots M}$ | G-dimensional vector of ambient feature counts | Ambient counts in cell *m* |
| $o_{m=1,\ldots M}$ | G-dimensional vector of final observation | Observed counts in cell *m* |

*Step 1, Generate native signals*

We first draw $\lambda$ from a Dirichlet distribution:

$$\lambda \sim Dirichlet\ (\xi)$$

The guide identities then can be sampled from Categorical distribution:

$$z_m \sim Categorical\ (G, \lambda)$$

$$y_m = Onehot(z_m)$$

CROP-seq libraries generally use an identical background construct, meaning all guides share an identical promoter. We therefore assume that per cell expression of native signals is drawn from a same distribution, independent of guide sequences. The expected native expression per cell can be calculated as:

$$\upsilon = \omega \times (1 - \varepsilon)$$

As there are technical dropout events, where there is a certain probability that molecules are not captured by sequencer. To reflect this, we draw the final native counts from a binomial distribution:

$$\sigma_m \sim Binomial\ (\upsilon, \varphi)$$

As we assume that each cell integrates only one guide in the case of CROP-seq, we can get the native expression vector for cell *m*:

$$n_m = \sigma_m \times y_m$$

*Step 2, Generate ambient signals*

Under ambient signal hypothesis, the ambient frequency α is correlated with the frequencies of native signals. And the native expression of guide is driven by the same promoter, so we have:

$$\alpha = \frac{\sum_M n_{mg}}{\sum_M \sum_G n_{mg}} = \lambda$$

Similarly, when considering the dropout event, we can draw actual ambient counts from a Binomial distribution:

$$\tau_m \sim Binomial\ (\omega \times \varepsilon, \varphi)$$

Then, the ambient signals per cell can be drawn from a Multinomial distribution:

$$a_m \sim Multinomial\ (\tau_m, \alpha)$$

*Step 3, Sum*

All together, we get the final observation,

$$o_m = n_m + a_m$$

2) Simulation of CITE-seq data

The definition of variables in this module is as follows.

| Variable | Data type | meaning |
|---|---|---|
| $M$ | Integer | Cell number |
| $G$ | Integer | Number of genes or features |
| $T$ | Integer | Number of cell types |
| $\xi$ | T-dimensional vector of positive real | Prior weight of cell types, 1 by default |
| λ | T-dimensional vector of probabilities, which must sum to 1 | Fraction of cell types |
| $\omega$ | Integer | Global sequencing depth, 8000 by default |
| $\psi_{t=1,...T}$ | Integer | The average of total native counts per cell in cell type *t* |
| $\varphi$ | Positive real between 0 and 1 | Capture rate, 0.7 by default |
| $\sigma_{m=1,...M}$ | Integer | Total native counts in cell *m* |
| $\delta$ | G-dimensional vector of positive real | Prior weight of genes (or features), 1/G by default to ensure sparsity |

| | | |
|---|---|---|
| $\beta_{t=1,...T}$ | G-dimensional vector of probabilities, which must sum to 1 | Distribution of native genes or features in cell type *t* |
| $n_{m=1, ...M}$ | G-dimensional vector of native feature counts | Native counts in cell *m* |
| α | G-dimensional vector of probabilities, which must sum to 1 | Distribution of ambient genes or features |
| $\rho$ | Positive real | The average of total ambient counts per cell |
| $z_{m=1, ...M}$ | Integer between 1 and T | Identity of cell type for cell *m* |
| $\tau_{m=1, ...M}$ | Integer | Total ambient counts in cell *m* |
| $a_{m=1, ...M}$ | G-dimensional vector of ambient feature counts | Ambient counts in cell *m* |
| $o_{m=1, ...M}$ | G-dimensional vector of final observation | Observed counts in cell *m* |

*Step 1, Generate native signals*

We first draw $\lambda$ from a Dirichlet distribution:

$$\lambda \sim Dirichlet\ (\xi)$$

The cell type identities then can be sampled from Categorical distribution:

$$z_m \sim Categorical\ (T, \lambda)$$

Different cell types generally have different total UMI counts (i.e., sequencing depth) in the case of CITE-seq protein counts, we therefore sample total native UMI counts per cell type from a discrete Uniform distribution:

$$\psi_t \sim U\ (\omega/2, \omega)$$

As there are technical dropout events, where there is a certain probability that molecules are not captured by sequencer. To reflect this, we draw the total native counts from a binomial distribution:

$$\sigma_m \sim Binomial\ (\psi_{t=z_m}, \varphi)$$

We next sample native expression frequencies ($\beta_t$) from a Dirichlet distribution:

$$\beta_t \sim Dirichlet_t\ (\delta)$$

Together, for cell *m*, the native expression at feature level can be drawn from multinomial distribution:

$$n_m \sim Multinomial\ (\sigma_m, \beta_{t=z_m})$$

*Step 2, Generate ambient signals*

Under ambient signal hypothesis, the ambient frequency α is correlated with the frequencies of native signals. So, we can calculate α using the following formula:

$$\alpha = \frac{\sum_M n_{mg}}{\sum_M \sum_G n_{mg}}$$

We assume that the total ambient counts are drawn from a discrete Uniform distribution:

$$\rho \sim U\ (0, \omega/2)$$

Similarly, when considering the dropout event, we can draw actual ambient counts from a Binomial distribution:

$$\tau_m \sim Binomial\ (\rho, \varphi)$$

Then, the ambient signals per cell can be drawn from a Multinomial distribution:

$$a_m \sim Multinomial\ (\tau_m, \alpha)$$

*Step 3, Sum*

All together, we get the final observation,

$$o_m = n_m + a_m$$
